# The IRE1 Inhibitor Kira6 Curtails The Inflammatory Trait Of Immunogenic Anticancer Treatments By Targeting Hsp60 Independent Of IRE1

**DOI:** 10.1101/2020.12.24.424330

**Authors:** Rufo Nicole, Korovesis Dimitris, Van Eygen Sofie, Rita Derua, Abhishek D Garg, Finotello Francesca, Vara-Perez Monica, Rožanc Jan, Dewaele Michael, Peter A de Witte, Leonidas G Alexopoulos, Sophie Janssens, Sinkkonen Lasse, Sauter Thomas, Steven HL Verhelst, Agostinis Patrizia

## Abstract

Cellular stress evoked by immunogenic anticancer treatments engaging the unfolded protein response (UPR) can elicit inflammation with conflicting therapeutic outcomes. To define cell-autonomous mechanisms coupling the UPR to molecular mediators of inflammation, we profiled the transcriptome of cancer cells responding to immunogenic or weakly immunogenic-treatments. Bioinformatics-driven pathway analysis indicated that immunogenic treatments instigated NF-κB/AP-1-inflammatory pathways, which were abolished by the IRE1α-kinase inhibitor KIRA6. Cell-free fractions of chemotherapy and KIRA6 co-treated cancer cells were deprived of pro-inflammatory/chemoattractant factors and failed to mobilize innate immune cells. Strikingly, these potent KIRA6 anti-inflammatory effects were found to be independent of IRE1α. Generation of a KIRA6-clickable photoaffinity probe, mass spectrometry and co-immunoprecipitation analysis identified cytosolic HSP60 as a KIRA6 off-target in the NF-κB pathway. In sum, our study unravels that inflammation evoked by immunogenic treatments is curtailed by KIRA6 independently of IRE1α and further suggests great caution in interpreting the anti-inflammatory action of IRE1α chemical inhibitors.

## INTRODUCTION

In response to anticancer treatments, stressed or dying cancer cells engage in a complex dialogue with the tumor microenvironment, which can ultimately stimulate or suppress inflammatory and immune responses. Various conventional or targeted therapies drive cancer cell-autonomous pathways leading to a form of immunostimulatory apoptosis, which is also called immunogenic cell death (ICD). Cancer cells responding to these immunogenic therapies enhance the microenvironmental availability of immunomodulatory or danger signals, which eventually favor adaptive immune responses specific for tumor-relevant antigens (Galluzzi et al. 2016). Scrutiny of the mechanistic underpinnings of ICD and *in vivo* studies have disclosed the pivotal role of the PERK/eIF2α-P axis of the unfolded protein response (UPR) following perturbation of ER homeostasis, as apical cancer cell-autonomous stress pathway orchestrating danger signaling (Krysko et al. 2012; Garg et al. 2012; Obeid et al. 2007). In contrast, the role of the IRE1α branch of the UPR during ICD remains insufficiently understood.

The UPR is also intimately involved in the activation of inflammatory responses, which classically involve NF-κB activation (Rufo, Garg, and Agostinis 2017). By phosphorylating eIF2α, PERK transiently attenuates global protein synthesis thereby favoring NF-κB activation of pro-inflammatory genes. IRE1α regulation of pro-inflammatory cytokine production can occur both via its RNase activity or its scaffolding role in the IRE1α-TRAF2-JNK pathway, eliciting NF-κB and AP-1 transcriptional pro-inflammatory programs (Urano et al. 2000; Hu et al. 2006). Recent studies in cancer (Lhomond et al. 2018; Logue et al. 2018; Pluquet et al. 2013) highlight that IRE1α induced expression of pro-inflammatory cytokines promotes the recruitment of pro-tumorigenic myeloid cells. Based on these findings pharmacological targeting of IRE1α has been suggested as a promising therapeutic avenue in different cancer settings.

However, while the critical role of danger signals has been validated in preclinical and in some clinical settings (Galluzzi et al. 2020), much less information is available on the molecular pathways controlling the pro-inflammatory outputs of the UPR elicited by immunogenic treatments. Moreover, whether a transcriptional inflammatory signature portrays a distinguished trait of immunogenic therapies has not been explored yet.

Several lines of evidence indicate that inflammation evoked by perturbations of ER homeostasis ignited by immunogenic therapies may play distinct role from danger signals (Sullivan et al. 2020) and facilitate autocrine/paracrine tissue regeneration responses with contrasting effects on disease progression. For example, in response to immunogenic chemotherapies, such as doxorubicin, CCL2 release by the stressed cancer cells boosts the initial phase of the immunogenic response by enabling the recruitment of antigen-presenting cells (Ma et al. 2014), but it can also evoke secondary inflammation, originating from both cancer cells and endothelial cells, which ultimately drives angiogenesis, metastasis and recurrence (Keklikoglou et al. 2019; Toste et al. 2016; Vyas, Laput, and Vyas 2014). Cancer vaccines generated by exposing cancer cells to immunogenic treatments, through the release of chemokines patterning cellular responses to pathogens, favor the recruitment of neutrophils at the site of vaccination, which enables *in vitro* killing of live cancer cells (Garg et al. 2017). However, in a tumor context, similar chemokines driving neutrophil infiltration into the tumor through the binding of their cognate CXCR1 or CXCR2 receptors, may favor tumor immunoescape by shielding cancer cells from cytotoxic T cells (Teijeira et al. 2020). Hence, depending on the context, sterile inflammatory responses elicited by stressed cancer cells may exert contrasting effects on the tumor microenvironment and on the overall output of immunogenic therapies. This creates an urgent need to delineate the cancer cell-autonomous molecular pathway responsible for the expression of these inflammatory factors.

In this study, we show that immunogenic treatments are hallmarked by a cancer cell-autonomous transcriptional footprint culminating in the NF-κB /AP-1 regulated expression of a common subset of pro-inflammatory chemokines. We reveal that the IRE1α kinase inhibitor KIRA6 curtails the pro-inflammatory output elicited by the stressed cancer cells exposed to immunogenic treatments. Strikingly, we found that the inhibitory effects of KIRA6 are completely independent of IRE1α and identified HSP60 as one of the off-target effectors of the KIRA6 anti-inflammatory action.

## RESULTS

### Immunogenic treatments are hallmarked by a pro-inflammatory transcriptional signature

We set out to investigate the transcriptional profile of cancer cells responding to mitoxantrone (MTX) or hypericin-based photodynamic therapy (Hyp-PDT), as prototypes of immunogenic treatments (Dudek et al. 2013). Cisplatin (CDDP) was chosen for comparison since it is considered a poorly immunogenic chemotherapeutic (Dudek et al. 2013). We first confirmed that these treatments induced similar kinetics of caspase-dependent cell death in the human melanoma A375 cell line (Figure 1 – figure supplement 1A), which we used in previous studies to characterize the involvement of ER stress induced inflammatory signals following melphalan chemotherapy (Dudek-Peric et al. 2015). Confirming previous reports (Sukkurwala et al. 2014; Michaud et al. 2014; Garg et al. 2012), MTX and Hyp-PDT elicited classical *in vitro* markers of immunogenic apoptosis, including surface exposed CRT (Figure 1 – figure supplement 1B), ATP release -particularly after Hyp-PDT (Figure 1 – figure supplement 1C) and the cytoplasmic redistribution of the nuclear HMGB1 prior to its passive release (Figure 1 – figure supplement 1D-E). In contrast, CDDP failed to induce these *in vitro* markers of immunogenic apoptosis (Figure 1 – figure supplement 1B-D).

We then interrogated the time dependent changes in the transcriptome of the treated A375 cells by bulk-RNA sequencing (RNAseq) analysis. We isolated the bulk-RNA of the treated cells, at 0 hr (untreated) and 4 hr (pre-apoptotic), 10 hr (early apoptotic) and 20 hr (late apoptotic) post-treatment, and compared it to their respective time-matched untreated controls (Figure 1 – figure supplement 1F). Differential gene expression analysis revealed that MTX and Hyp-PDT significantly upregulated the transcription of several genes as early as 4 hr post-treatment, whereas responses to CDDP treatment occurred mainly at the later time points and were associated with a predominant transcriptome downregulation with respect to the untreated controls (Figure 1 – figure supplement 1G). We then performed principal component analysis (PCA) on the top 10% most variable genes considering rlog-normalized RNAseq counts. Bidimensional PCA output showed that MTX and Hyp-PDT treatments clearly separate on the second dimension from CDDP, which accounted for 30% of the total variance (Figure 1A). To explore if genes discriminating between these treatments could be assigned to specific pathways, we ran a Gene Set Enrichment Analysis (GSEA) (Subramanian et al. 2005) using the WikiPathway Cancer gene set and weighing the genes for their contribution to the principal component 1 (PC1) and PC2 (Figure 1B). Pathways clustering the two genotoxic agents CDDP/MTX in the PC1 were as expected enriched in the “DNA-damage response” gene set (Figure 1B). Instead, “chemokine signaling pathway” emerged as the predominantly enriched gene signature from the second dimension (PC2) which clustered MTX and Hyp-PDT (Figure 1B). Gene Ontology (GO) analysis performed on the totality of significantly upregulated genes pinpointed “inflammatory response/immune responses” and “innate immune responses” as the most significant GO terms belonging to biological processes, and “extracellular space” as most significant term belonging to cellular compartment, almost exclusively in association with immunogenic treatments (Figure 1C). This GO analysis highlighted that differential expression of genes belonging to these biological processes were upregulated at the pre-apoptotic phase (i.e. 4 hr after treatment) and further remained elevated through the full time course (Figure 1C). Genes regulated by the PERK- and IRE1α-mediated UPR responses were particularly sustained after Hyp-PDT (Figure 1C). MTX elevated transiently PERK-regulated genes while CDDP failed to evoke UPR-regulated genes (Figure 1C). Immunoblot analysis further substantiated that Hyp-PDT - a primarily ER-targeted ROS generating treatment (Garg et al. 2012; Verfaillie et al. 2012; Agostinis et al. 2011)- promoted rapid phosphorylation of IRE1α and elevated the expression of the spliced form of *XBP1* (*sXBP1,* figure supplement 1H), resulting from the activation of the IRE1α RNase activity. Hyp-PDT induced eIF2α phosphorylation (a main PERK downstream effector) (Figure 1D) and elicited markers of terminal UPR, including ATF4, CHOP and its downstream target DR5 (TRAIL-R2) (Figure 1E). MTX significantly stimulated the phosphorylation of IRE1α (Figure 1E) but failed to elevate the expression of s*XBP1 (*Figure 1 – figure supplement 1H), suggesting its inability to promote IRE1α RNase activity. MTX elicited eIF2α phosphorylation, a signature of the integrated stress response (ISR) (Bezu et al. 2018), and increased the expression of DR5 at later time points (Figure 1E). In contrast, CDDP failed to induce these ER stress markers to a significant extent (Figure 1D,E).

**Figure 1.**
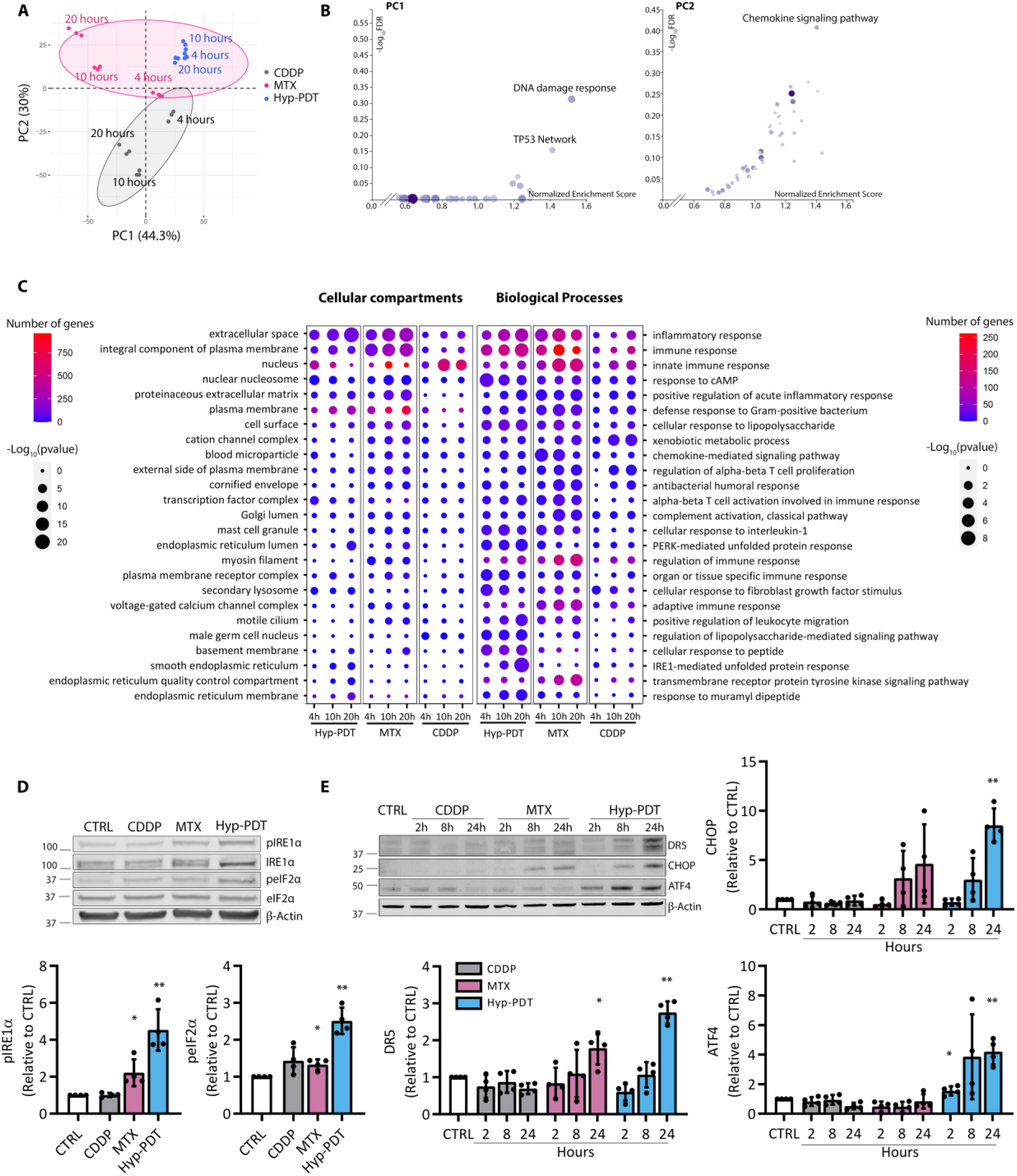
Immunogenic treatments are hallmarked by a pro-inflammatory transcriptional program. **(A)** Principal component analysis (PCA) performed on the rlog-normalized RNAseq counts from A375 cells at 4 hr, 10 hr and 20 hr after treatment with CDDP, MTX or Hyp-PDT. Three independent biological replicates are represented for each time point and treatment. Ellipses draw 90% confidence area for immunogenic (pink) and non-immunogenic (CDDP, grey) treatments. **(B)** Gene set enrichment analysis (GSEA) of Wikipathways Cancer showing the ‒Log10(FDR) and Normalized enrichment scores of the gene sets positively associated to input gene lists. **(C)** Results of the Gene ontology (GO) analysis based on genes significantly upregulated in the RNAseq dataset in samples treated with CDDP, MTX or Hyp-PDT compared to time-matched untreated control. Top 25 most variable GO terms relative to biological process and cellular compartment are represented. Dot sizes and colors represent ‒log10(p-value) of the enrichment of each term in the different samples and number of genes, respectively. **(D)** Representative western blot and relative quantifications for early markers (phosphorylated IRE1α (pIRE1α) and peIF2α, 2 hr post-treatment) and **(E)** time course of late markers (ATF4, DR5 and CHOP) underlining UPR induction after treatment with CDDP, MTX and Hyp-PDT. β-actin was used as normalization parameter; peIF2α is normalized over total levels of eIF2α. Values represent mean ± SD of the fold change over control. Data is analyzed by one sample t-test against hypothetical value control set to 1. *p<0.05, **p<0.01.

Thus, immunogenic treatments engaging the UPR/ISR evoke a distinct transcriptional signature hallmarked by the expression of a subset of pro-inflammatory transcripts.

### Transcriptional upregulation of a subset of chemokines is a hallmark of immunogenic treatments

To further unravel the transcriptional signature differentiating immunogenic treatments from CDDP, we focused on pro-inflammatory entities, which were consistently upregulated by both MTX and Hyp-PDT, while concomitantly down-regulated in response to CDDP. This analysis identified *CXCL8*, *CXCL3* and *CXCL2* as the subset of chemokines differentially co-upregulated in response to immunogenic treatments, while simultaneously co-repressed by CDDP (Figure 2A).

**Figure 2.**
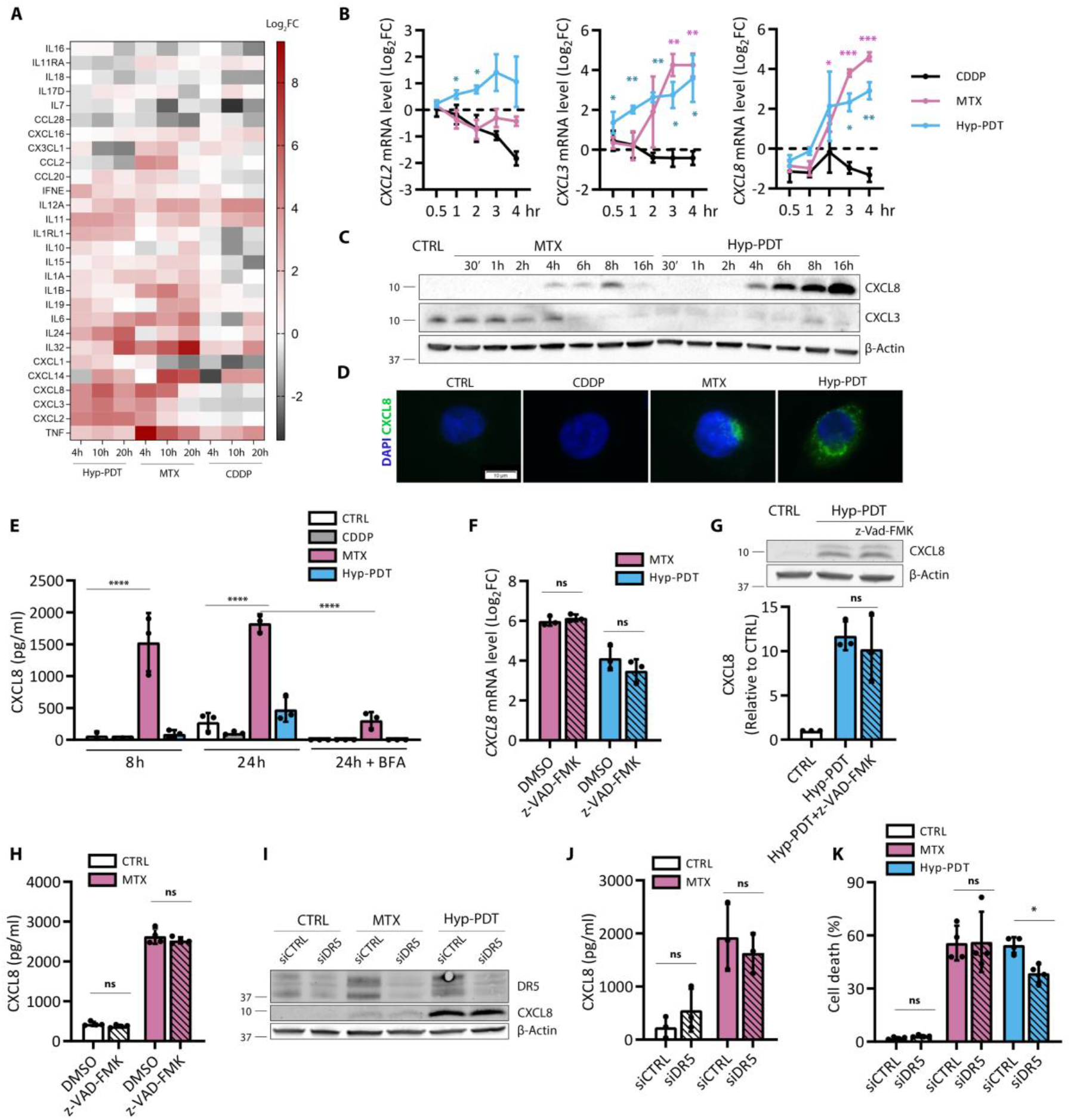
Immunogenic treatments elicit chemokine production independent of cell death. **(A)** Heatmap obtained from RNAseq data representing the log2fold changes of cytokine and chemokines genes upon treatment with CDDP, MTX and Hyp-PDT compared to time-matched untreated controls. **(B)** Chemokine transcription was evaluated at short time points after treatment with CDDP, MTX and Hyp-PDT with a time course analysis performed by qPCR. Data is normalized on 18s housekeeping gene and expressed as log2(fold change) compared to time-matched untreated control. Asterisks are color matched to the treatment. **(C)** Representative western blot of intracellular CXCL8 accumulation and CXCL3 depletion at the indicated time-points upon treatment with MTX and Hyp-PDT. **(D)** Representative fluorescence microscopy pictures visualizing the intracellular accumulation of CXCL8 (green) in control (CTRL) condition or 8h after treatment with CDDP, MTX or Hyp-PDT. Nuclei were counterstained with DAPI (blue). Scale bar: 10 μM **(E)** CXCL8 secretion was measured by ELISA in conditioned medium at 8 hr and 24 hr after treatment with CDDP, MTX or Hyp-PDT. CXCL8 secretion at 24 hr after treatment was blocked by inhibiting anterograde transport with Brefeldin A (BFA, 50 ng/ml). **(F-H)** Evaluation of the impact of cell death inhibition by z-VAD-FMK (50 μM) on CXCL8 transcription (F), intracellular protein accumulation (G) and secretion (H) after treatment with Hyp-PDT and/or MTX as assessed by qPCR, Western Blot and ELISA, respectively. Values are represented as Log2(fold change) and fold change over untreated control in (F) and (G), respectively. **(I)** Impact of siDR5 on CXCL8 protein production or **(J)** secretion, measured by Western Blot and ELISA, respectively. **(K)** Cell death was assessed in siRNA-mediated DR5 knock-down compared to scrambled siRNA (siCTRL) 24 hr after treatment with MTX and Hyp-PDT. In all Western Blots β-actin was used as loading control. In all graphs values are presented as mean ± SD. Data is analyzed by Two-way ANOVA in all the graphs except for one sample t-test in (B) and Student’s t-test in (G), *p<0.05,**p<0.01, ***p<0.001, ****p<0.0001, ns=not significant.

We then validated the RNAseq data by kinetic qPCR kinetic analysis. Matching the RNAseq data, *CXCL3* and *CXCL8* were rapidly induced in response to MTX and Hyp-PDT, whereas the upregulation of *CXCL2* was less pronounced and restricted to Hyp-PDT (Figure 2B). The expression of these chemokines was downregulated upon treatment with CDDP (Figure 2A, B).

We next focused on the production of CXCL3 and CXCL8 as these chemokines were most significantly induced by both MTX and Hyp-PDT and are generally characterized by their ability to attract myeloid cells, like neutrophils, and to exert relevant autocrine and paracrine functions (Liu et al. 2016).

CXCL3 protein levels were expressed at baseline but decreased following the treatments (Figure 2C). Bortezomib, an inhibitor of the proteasome (Figure 2 – figure supplement 1A), restored CXCL3 protein levels, suggesting that following its intracellular accumulation this chemokine is subjected to a fast degradation. On the contrary, CXCL8 levels were undetectable in untreated cells but increased rapidly starting from 4 hr after exposure to MTX or Hyp-PDT, as indicated by immunoblotting and immunostaining (Figure 2C, D). In response to MTX intracellular accumulation CXCL8 was accompanied by its secretion, which remained sustained through the time (Figure 2E). The release of CXCL8 was blocked by the secretory pathway inhibitor Brefeldin A indicating that it occurred via canonical anterograde transport (Figure 2E, Figure 2 – figure supplement 1B). In contrast, after Hyp-PDT, CXCL8 accumulated predominantly intracellularly and was poorly or not significantly released at later time points (Figure 2E), in line with the reported reduced secretory ability of Hyp-PDT stressed cells (Gomes‐da‐Silva et al. 2018; Garg et al. 2012; Dudek-Peric et al. 2015).

We then focused our mechanistic analysis on CXCL8 as prototype of pro-inflammatory chemokine induced by both immunogenic treatments in stressed/dying cancer cells.

We then evaluated whether global blockade of caspases had an effect on CXCL8 induction and secretion after immunogenic treatments. Blocking apoptosis by the pan-caspase inhibitor z-VAD-FMK (Figure 1 – figure supplement 1A) had no effect on the induction of CXCL8 at the RNA level (Figure 2F), intracellular protein accumulation (Figure 2G) and on its release after MTX (Figure 2H). Thus increasing the fraction of surviving cells did not affect CXCL8 production after immunogenic treatments.

In response to classical ER stress inducing agents like thapsigargin or tunicamycin, or following treatment with microtubule disruptors (taxanes) ligand independent, PERK-mediated and CHOP-induced upregulation of DR5 (TRAIL-R2) triggers apoptosis concurrent to NF-κB driven inflammation (Sullivan et al. 2020; Lam et al. 2018). After MTX or Hyp-PDT, the upregulation of CXCL8 at the mRNA (Figure 2B) and protein levels (Figure 2C) occurred at the pre-apoptotic phases (e.g. as early as 2-4 hr post-treatment). In contrast, the upregulation of CHOP and DR5 (Figure 1F) was a late event associated to apoptotic cell death (i.e. 24 hr post-treatment). In spite of the unrelated kinetics, we next tested the effect of DR5 silencing on CXCL8 production and cell death after immunogenic treatments. The siRNA-mediated silencing of DR5 (about 75%, Figure 2I) did not affect CXCL8 production after treatment with neither Hyp-PDT nor MTX (Figure 2I, J). Downregulation of DR5 expression or pharmacological blockade of PERK (Figure 2 – figure supplement 1C) reduced to some extent cell death after Hyp-PDT (Figure 2K). This is consistent with the pro-apoptotic role of PERK after this ROS-induced ER stress treatment, which involves the canonical PERK-CHOP axis and a kinase-independent role of PERK at the ER-mitochondria contact sites (Verfaillie et al. 2012).

Collectively, these results show that CXCL8 production following immunogenic treatments occurs as a result of a DR5- and caspase-independent pre-mortem induced stress response.

### CXCL8 induction following immunogenic treatments requires NF-κB and cJUN

The finding that CXCL8 production occurred through DR5- and caspase-independent mechanisms suggests that immunogenic drugs engage other circuits leading to NF-κB activation following ER stress/eIF2α phosphorylation or evoke the activation of other transcription factors (TFs). We then queried three different bioinformatics tools (IPA, iRegulon and GATHER) to predict putative transcription factors controlling shared Hyp-PDT- and MTX-induced transcripts belonging to the GO term “extracellular space”. Only two TFs were consistently present in the top 10 by all three tools, namely the AP-1 member cJUN and NF-κB (Figure 3A). In the canonical NF-κB pathway, activation of the inhibitor of the IκB kinase (IKK) complex drives phosphorylation-mediated IκBα degradation, thus licensing NF-κB activation (Israël 2010).

**Figure 3.**
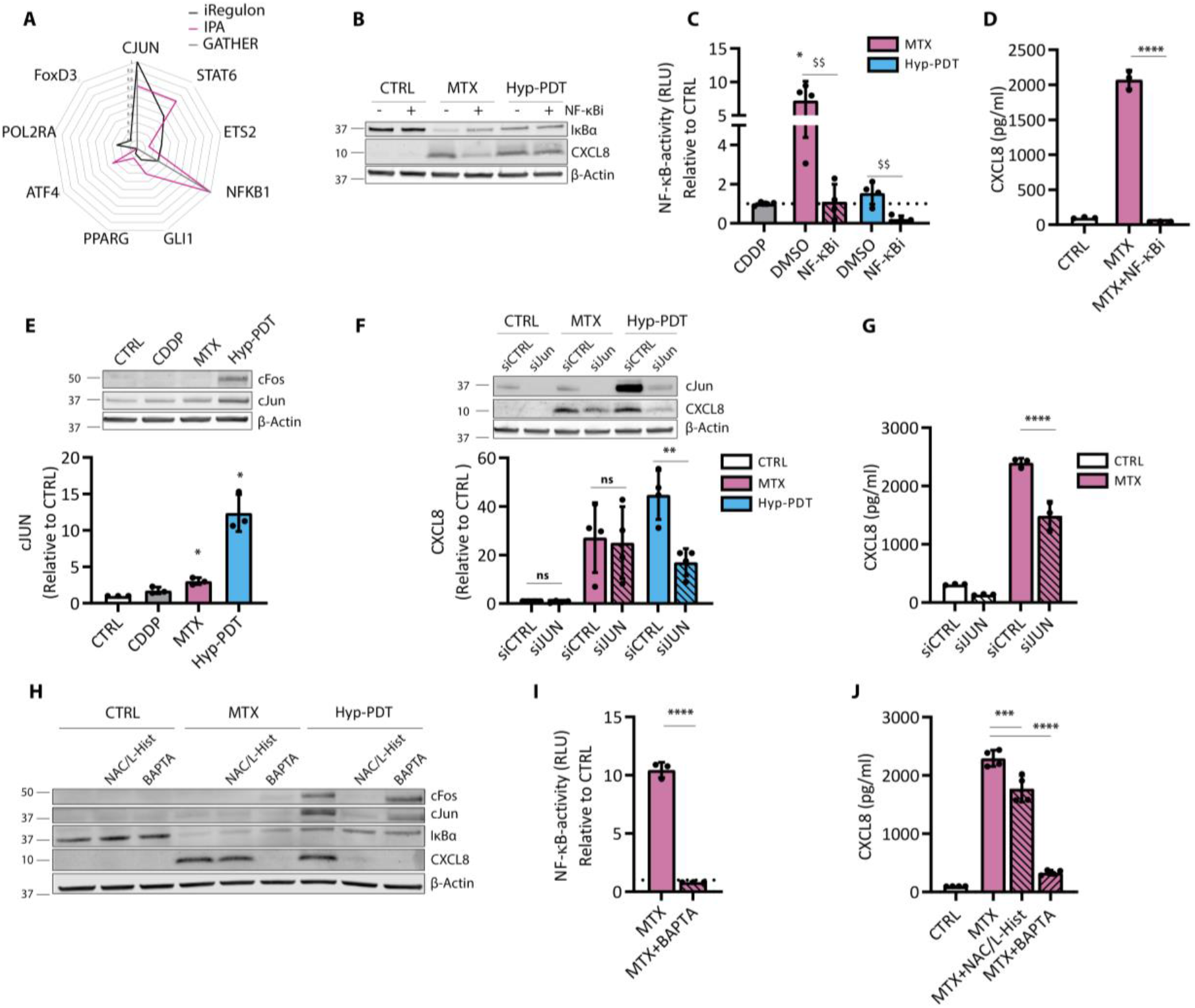
CXCL8 production is mediated by cJUN and NF-κB. **(A)** In silico prediction of transcription factors involved in the regulation of genes belonging to the “extracellular space” GO term significantly upregulated by both MTX and Hyp-PDT. The transcription factors reported are derived from iRegulon prediction, and independent scores of the same transcription factors by IPA and Gather are provided. Data is reported after min-max normalization. **(B)** Representative Western Blot showing the impact of BAY11-7082 (10 μM) on degradation of IκBα and CXCL8 intracellular protein accumulation in basal condition (CTRL) and 4 hr after treatment with MTX or Hyp-PDT. **(C)** NF-κB activity measured by luciferase assay in A375 cells stably expressing the reporter 4 hr after treatment with CDDP, MTX and Hyp-PDT in the presence or absence of BAY11-7082 (10 μM). Data is expressed as fold change compared to untreated control, whose reference value is indicated with a dotted line. **(D)** CXCL8 secretion measured by ELISA in conditioned medium from A375 cells with or without co-incubation with BAY11-7082 (10 μM) 24 hr after treatment with MTX. **(E)** Representative Western Blot of intracellular levels of cFOS and cJUN and quantification 4 hr after treatment with CDDP, MTX or Hyp-PDT. Data is expressed as fold change over untreated control. **(F)** Representative Western Blot and quantification of the impact of siRNA mediated knock-down of cJUN (siJUN) with respect to scramble siRNA (siCTRL) on intracellular CXCL8 accumulation 4 hr after treatment with MTX and Hyp-PDT. Data is expressed as fold change over control incubated with siCTRL. **(G)** CXCL8 secretion was measured by ELISA in the conditioned medium from A375 cells with siRNA mediated cJUN knock-down 24 hr after treatment with MTX. **(H)** Representative Western Blot evaluating the impact of ROS inhibitor N-Acetyl cysteine (NAC, 5 mM))/L-Histidine (25 mM) and cell permeable calcium chelator BAPTA-AM (10 μM) on the upregulation of cJUN and cFOS, degradation of IκBα and CXCL8 production 4 hr after treatment with MTX or Hyp-PDT. **(I)** NF-κB activity measured by luciferase assay in A375 cells stably expressing the reporter 4 hr after treatment with MTX in presence or absence of BAPTA-AM (10 μM). Data is expressed as fold change compared to untreated control, whose reference value is indicated with a dotted line. **(J)** CXCL8 secretion measured by ELISA in conditioned medium from A375 cells with or without co-incubation with NAC (5 mM)/L-Histidine (25 mM) or BAPTA-AM (10 μM) 24 hr after treatment with MTX. In all Western Blots β-actin was used as loading control. In all graphs values are presented as mean ± SD. Data is analyzed by one-sample t-test in (C) and (E), Student’s t-test in (C) with respect to treatment, (D) and (I), Two-way ANOVA in (F) and (G) and One-way ANOVA in (J). *p<0.05, **p<0.01, ***p<0.001, ****p<0.0001, $$p<0.001, ns=not significant.

In line with this, MTX caused degradation of IκBα (Figure 3B), which was accompanied by NF-κB activation, as measured by a luciferase reporter assay (Figure 3C). MTX-stimulated CXCL8 production (Figure 3B, D) and NF-κB activation (Figure 3C) were blunted by the IKK inhibitor Bay 11-7082, which blocked IκBα degradation (Figure 3B). Hyp-PDT failed to stimulate NF-κB signaling (Figure 3B,C), as reported in previous studies (Hendrickx et al. 2003) using different human cancer cell lines, and accordingly CXCL8 induction by this immunogenic treatment was not affected by NF-κB inhibition (Figure 3 – figure supplement 1A). Instead, Hyp-PDT induced a robust upregulation of the main AP-1 members cJUN and cFOS (Assefa et al. 1999), while MTX partially, albeit significantly affected cJUN expression (Figure 3E). Silencing cJUN decreased intracellular expression of CXCL8 in response to Hyp-PDT (Figure 3F) and reduced CXCL8 secretion after MTX (Figure 3G). The complete inhibition of CXCL8 production by the NF-κB inhibitor (Figure 3C) suggests that following MTX, NF-κB operates as a dominant pro-inflammatory TF, which facilitates cJUN mediated transcription as reported in previous studies (Fujioka et al. 2004). Accordingly, we observed that NF-κB inhibition significantly decreased cJUN upregulation in MTX-treated cells (Figure 3 – figure supplement 1B). CDDP failed to stimulate these TFs (Figure 3C, E) in agreement with its inability to induce CXCL8 and the pattern of cytokines associated to immunogenic treatments (Figure 2A, B).

Interestingly, impairing elevation of intracellular Ca^2+^ or the generation of ROS, two apical elements of the cascade leading to immunogenic apoptosis (Panaretakis et al. 2009; Garg et al. 2012; Martins et al. 2011), significantly curtailed both NF-κB and AP-1 mediated CXCL8 production (Figure 3H-J). Together these results show that the pro-inflammatory output elicited by immunogenic treatments is mediated by NF-κB and AP-1 signaling.

### Pharmacological inhibition of IRE1α kinase activity curtails pro-inflammatory responses after immunogenic chemotherapy

Upon detection of unfolded proteins in the ER lumen by their luminal domains, PERK and IRE1α oligomerize and trans-autophosphorylate via their cytoplasmic Ser/Thr kinase domain. This apical event leads to PERK phosphorylation of eIF2α and to the allosteric activation of the IRE1α RNase activity (Hetz, Chevet, and Oakes 2015).

Both PERK and IRE1α pathways mediate NF-κB and AP-1 driven pro-inflammatory cytokine production (Hetz, Zhang, and Kaufman 2020) and pharmacological inhibitors of these ER stress sensors have been used *in vivo* to ameliorate the pathological output of the UPR in cancer and other inflammatory diseases (Grandjean and Wiseman 2020).

We therefore tested the effects of PERK and IRE1α chemical inhibitors on CXCL8 production after immunogenic treatments. The inhibition of PERK had only a marginal (albeit significant) effect on CXCL8 production by MTX (Figure 4A) or no effect after Hyp-PDT (Figure 4 – figure supplement 1A). In contrast, blocking the autophosphorylation of IRE1α by KIRA6 -a prototype compound of the small molecules Kinase Inhibiting RNase Attenuators (Ghosh et al. 2014) (Figure 4 – figure supplement 1B)- curtailed CXCL8 production in response to both MTX and Hyp-PDT (Figure 4A, Figure 4 – figure supplement 1A).

**Figure 4.**
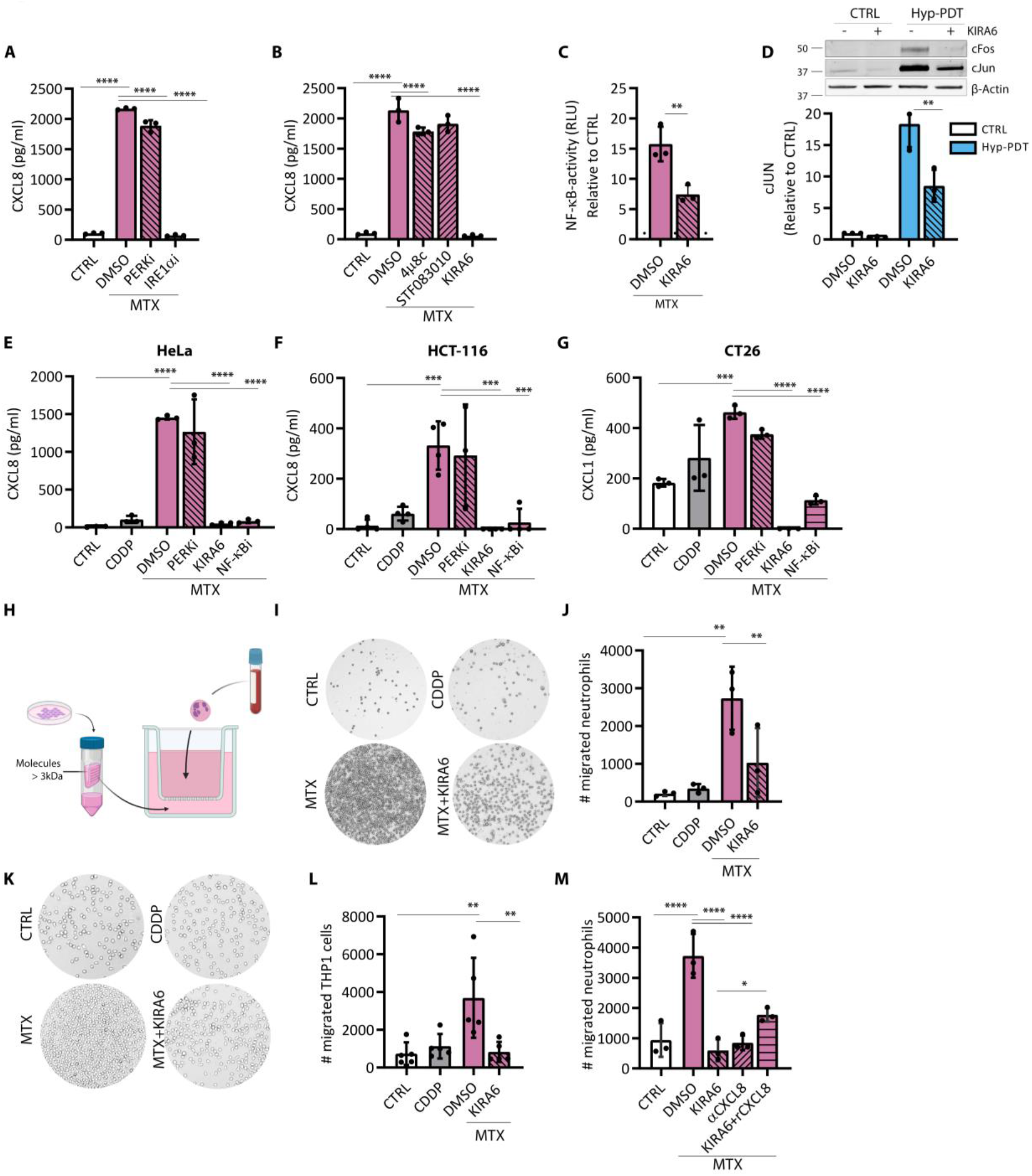
The IRE1α kinase inhibitor KIRA6 blunts CXCL8 production and neutrophil recruitment after immunogenic treatment. **(A)** CXCL8 secretion was measured by ELISA 24 hr after treatment with MTX in conditioned medium of A375 cells in coincubation with the PERK inhibitor GSK2606414 (1 μM) or the IRE1α kinase inhibitor KIRA6 (1 μM). **(B)** CXCL8 secretion was measured by ELISA 24 hr after treatment with MTX in conditioned medium of A375 cells in coincubation with the IRE1α RNAse inhibitors 4μ8C (100μM), STF-083010 (50μM) or IRE1α kinase inhibitor KIRA6 (1 μM). **(C)** NF-κB activity measured by luciferase assay in A375 cells stably expressing the reporter 4 hr after treatment with MTX in the presence or absence of the IRE1α kinase inhibitor KIRA6 (1 μM). Data is expressed as fold change compared to untreated control, whose reference value is indicated with a dotted line. **(D)** Impact of IRE1α kinase inhibitor KIRA6 (1 μM) on intracellular levels of cJUN and cFOS in basal conditions (CTRL) and 4 hr after treatment with Hyp-PDT. β-actin was used as loading control. **(E-F)** CXCL8 secretion was measured by ELISA 24 hr after treatment with MTX in conditioned medium of HeLa and HCT-116 cells in coincubation with the PERK inhibitor GSK2606414 (1 μM), the IRE1α kinase inhibitor KIRA6 (1 μM) or NF-κB inhibitor BAY11-7082 (10 μM). **(G)** CXCL1 secretion was measured by ELISA 24 hr after treatment with MTX in conditioned medium of murine CT26 cells in coincubation with PERK inhibitor GSK2606414 (1 μM), IRE1α kinase inhibitor KIRA6 (1 μM) and NF-κB inhibitor BAY11-7082 (10 μM). **(H)** Scheme of the transwell migration experimental setup. **(I-J)** Representative pictures and relative quantification of the transwell migration assay of human neutrophils exposed for 2 hr to conditioned medium of A375 cells treated for 24 hr with MTX in the presence or absence of the IRE1α kinase inhibitor KIRA6 (1 μM). **(K-L)** Representative pictures and relative quantification of the transwell migration assay of human macrophage-like THP1 cell line exposed for 4 hr to conditioned medium A375 cells treated for 24 hr with MTX in the presence or absence of IRE1α kinase inhibitor KIRA6 (1 μM). **(M)** Neutrophil’s migration was assessed after neutralization of CXCL8 secreted upon treatment with MTX with a αCXCL8 neutralizing antibody (0.5 μg/ml) or after addition of recombinant CXCL8 in MTX+KIRA6 conditioned medium at a concentration of 10 ng/ml. In all graphs values are presented as mean ± SD. Data is analyzed by One-way ANOVA in all the graphs except for Student’s t-test in (C). *p<0.05, **p<0.01, ***p<0.001, ****p<0.0001.

The imidazopyrazine KIRA6 compound interacts with the ATP binding site of IRE1α. This causes the inhibition of IRE1α autophosphorylation (Figure 4 – figure supplement 1B) and oligomerization, and consequently allosteric inhibition of the RNase activity of IRE1α (Harnoss et al. 2019) (Figure 4 – figure supplement 1C). IRE1α phosphorylation, however, can signal to the activation of NF-κB and AP-1 through the recruitment of scaffolding proteins, like TRAF2 and Nck, independent of its RNAse activity and in response to stress signals elevating intracellular Ca^2+^ and ROS (Adams et al. 2019; Nguyên et al. 2004; Urano et al. 2000; Son et al. 2014; Tufanli et al. 2017). To further elucidate the mechanistic underpinnings of the pro-inflammatory pathway controlled by IRE1α, we then evaluated if inhibitors of its RNase activity phenocopied the effects of KIRA6. Blocking IRE1α RNase activity by the chemical inhibitors 4μ8c or STF083010 (Figure 4 – figure supplement 1C) had no or only marginal effects on the production of CXLC8 following MTX (Figure 4B), in line with its inability to stimulate expression of *sXBP1* (Figure 1 – figure supplement 1G), or Hyp-PDT (Figure 4 – figure supplement 1D) while KIRA6 abrogated it (Figure 4B, Figure 4 – figure supplement 1D). These results suggested that IRE1α kinase activity is necessary to ignite the pro-inflammatory program, possibly through its scaffolding ability.

KIRA6 significantly reduced both NF-κB activity following MTX (Figure 4C) and AP-1 induction following Hyp-PDT (Figure 4D). We then explored if the KIRA6 inhibitory effects on CXCL8 production after immunogenic therapy were observed in other cancer cell lines of human and murine origin. Because Hyp-PDT treated cells failed to secrete chemokines, we treated these cancer cells with MTX and measured CXCL8 by ELISA. Similarly to what found in the A375 cells, only the blockade of NF-κB by Bay 11-7082 and of the IRE1α kinase by KIRA6 abolished MTX induced CXCL8 secretion by the human HeLa (Figure 4E) and HCT116 (Figure 4F) cells, and CXCL1 (a murine ortholog of CXCL8) secretion by the murine CT26 cells (Figure 4G) to a notably similar degree. We then assessed a wider panel of cytokines released by the A375 cells in response to MTX by multiplexed ELISA. We observed a general inhibition of NF-κB driven inflammation by KIRA6 (Figure 4 – figure supplement 1E), with the exception of an interferon-inducible CXCL10 (Metzemaekers et al. 2018) (Figure 4 – figure supplement 1E). This suggests that KIRA6 does not interfere with the production of all chemokines, but it inhibits mainly pathways leading to NF-κB-induced pro-inflammatory targets.

Since CXCL8 is the most potent human prototype of neutrophil-attracting chemokine belonging to the ELR^+^ CXC chemokine family (Nauseef and Borregaard 2014; Russo et al. 2014), we then evaluated the effects of KIRA6 on the ability of the cell-free supernatants of MTX-treated A375 cells to recruit human neutrophils. To exclude direct effects of the drug and chemical inhibitor on neutrophils, we processed the conditioned media to extensive rounds of sieving to retain only components heavier than 3 kDa, thus eliminating all small molecules and drugs while preserving chemokines (typically in the range of 7-15 kDa) (Figure 4H). The cell-free fraction from untreated or CDDP treated cells failed to attract human neutrophils isolated from peripheral blood of healthy donors in the transmigration assay (Figure 4H, I). In contrast, the conditioned media of the MTX treated cancer cells robustly stimulated the recruitment of neutrophils (Figure 4I,J), which was significantly blocked when performing the MTX treatment of cancer cells in the presence of KIRA6. Similar results were obtained by assessing the transmigration capacity of the human monocyte-like cell line THP1 (Figure 4K, L). Antibody-based neutralization of CXCL8 blocked neutrophils chemotaxis elicited by the conditioned media of MTX-treated cancer cells, to the same extent of that of the untreated cells or MTX-treated cells in the presence of KIRA6 (Figure 4M). Opposite to that, exogenous addition of recombinant CXCL8 to the KIRA6-conditioned medium significantly recovered neutrophil transmigration ability, albeit not completely (Figure 4M). This suggests that recombinant CXCL8 may not fully mimic the native conformation of the secreted CXCL8 (Mortier et al. 2011; Vacchini et al. 2018) or that internalization of CXCR1/CXCR2 receptors (Stone et al. 2017) dampened neutrophil migration in our assay. Taken together these results suggest that inhibition of IRE1α kinase activity by KIRA6 blunts the signal leading to the pro-inflammatory trait induced by immunogenic treatments, independently of the cancer cell type.

### The anti-inflammatory effects of KIRA6 are independent of IRE1

Inhibitors of the kinase site of IRE1α are known to either increase (Feldman et al. 2016) or decrease RNase activity (Maly and Papa 2014), by promoting or preventing IRE1α oligomerization, respectively (Carlesso et al. 2019). KIRA6 belongs to the latter class of ATP-competitive ligands, which allosterically inhibit IRE1α RNase (figure supplement 4C).

To validate the effects of KIRA6 in suppressing the inflammatory output of immunogenic treatments we next generated IRE1α knockout (KO) A375 cells through CRISPR/Cas9 gene editing. Efficient IRE1α KO was assessed by lack of detectable IRE1α protein and absent XBP1s cleavage (Figure 5A, Figure 5 – figure supplement 1A). To our surprise, IRE1α^−/−^ A375 cells were as proficient as their IRE1α^+/+^ counterparts in promoting CXCL8 production following treatment with either MTX or Hyp-PDT (Figure 5A, B). Moreover, KIRA6 was still able to block CXCL8 levels in IRE1α^−/−^ A375 cells exposed to MTX or Hyp-PDT (Figure 5A, B). Similar results were obtained when IRE1α was downregulated by siRNA treatment in A375 cells (Figure 5 – figure supplement 1B) or in IRE1α KO MEFs (Figure 5 – figure supplement 1C). Together these results indicated a hitherto unknown off-target effector of the anti-inflammatory activity of KIRA6 following immunogenic treatments.

**Figure 5.**
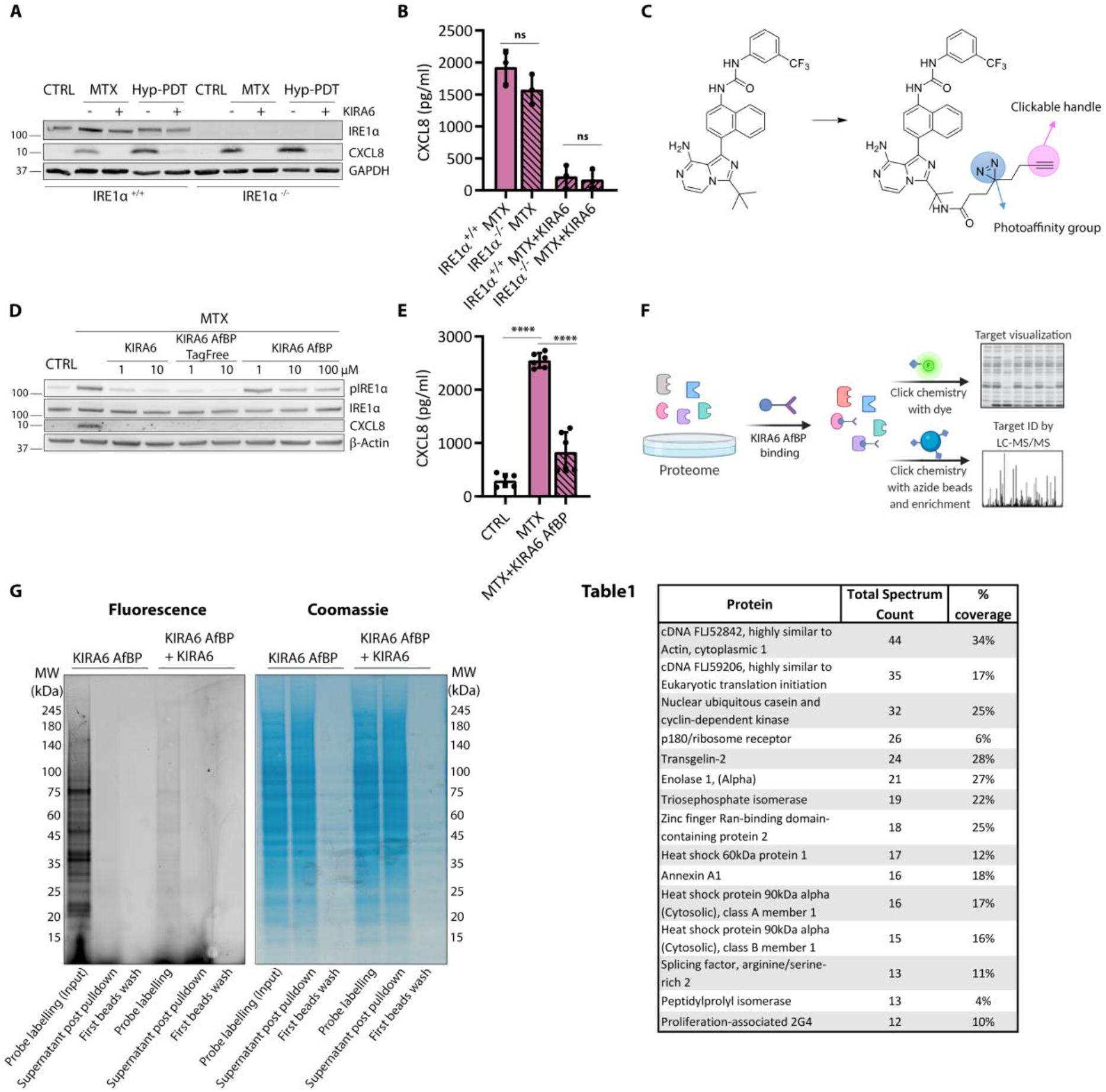
KIRA6 modulates CXCL8 production in an IRE1α independent pathway. **(A)** Representative Western Blot showing KIRA6 (1 μM) mediated blocking of CXCL8 production and intracellular accumulation 4 hr after treatment with MTX and Hyp-PDT in both IRE1α^−/−^ or IRE1α^+/+^ A375 cells. **(B)** CXCL8 secretion was measured by ELISA 24 hr after treatment with MTX in conditioned medium of IRE1α^−/−^ or IRE1α^+/+^ A375 cells with or without incubation with IRE1α kinase inhibitor KIRA6 (1 μM). **(C)** Molecular structure of the modified KIRA6 (KIRA6 Affinity-based Probe, KIRA6 AfBP) used for off-target protein identification. **(D)** Representative Western Blot comparing the ability of KIRA6 to KIRA6 AfBP and the intermediate tag-free KIRA6 AfBP (without photoaffinity group and clickable handle) in inhibition of CXCL8 production and IRE1α phosphorylation 4 hr after treatment with MTX in A375 cells at the indicated concentrations. **(E)** CXCL8 secretion was measured by ELISA 24 hr after treatment with MTX in conditioned medium of A375 cells with or without coincubation with KIRA6 AfPB (10 μM). **(F)** Scheme of the experimental workflow used to identify the KIRA6 off-targets by using the modified KIRA6 AfBP. **(G)** Representative gel assessing the efficacy of protein off-target identification workflow in 1 mg/ml protein lysates of A375 cells by reacting the photoaffinity-labeled lysates or azide bead supernatants with TAMRA-azide, after which SDS-PAGE and fluorescence scanning is performed. Lane 1 shows multiple fluorescent bands indicating KIRA6 AfBP (10 μM) labelled proteins. Lane 2 shows selective depletion of KIRA6 AfBP labeled proteins following azide beads pulldown indicated by decrease of fluorescent signal but a comparable amount of proteins (Coomassie). Lane 3 shows efficient removal of the most abundant aspecific proteins (Coomassie) but absence of removal of labeled proteins (fluorescence). Lane 4 shows that KIRA6 AfBP protein binding is out competed by coincubation with KIRA6 (10 μM) suggesting overlapping targets. (**Table 1)** Top 15 off-target candidates obtained by LC-MS/MS scored by total spectral counts. In the graphs values are presented as mean ± SD and analyzed by Two-way ANOVA in (B) and One-way ANOVA in (E), ****p<0.0001, ns= not significant.

In order to identify the off-target effector of KIRA6, we utilized a novel photoaffinity and clickable KIRA6 probe (KIRA6 AfBP) (Figure 5C), which we recently synthesized (Korovesis et al. 2020) by incorporating a diazirine photoreactive group and an alkyne biorthogonal tagging moiety onto the KIRA6 imidazopyrazine scaffold. The KIRA6 AfBP allowed the pull down of the covalently bound (off-)targets and their subsequent identification by mass spectrometry (MS).

We first tested KIRA6 AfBP for its ability to suppress CXCL8 production in A375 cells treated with MTX. KIRA6 AfBP displayed comparable inhibitory efficiency, when compared to KIRA6, in decreasing CXCL8 levels from MTX-treated cells (Figure 5D, E). Interestingly, the inclusion of the tag (i.e. photoreactive group+clickable handle) prevented the binding to IRE1α in live cells, since phosphorylation of IRE1α was still detectable when using KIRA6 AfBP, whereas it was suppressed by the tag-free KIRA6 AfBP precursor (Figure 5D). This suggests that in the A375 cells, KIRA AfBP is only able to bind to the off-target partner(s) of KIRA6, while leaving IRE1α activity intact. Thus, while KIRA AfBP preserved IRE1α phosphorylation, it still repressed CXCL8 production, again underscoring the IRE1α-independent effect on the pro-inflammatory responses initiated by immunogenic treatments.

We then treated A375 cells with KIRA AfBP, followed by photocrosslinking to covalently modify protein targets. Subsequently, cells were lysed and probe-modified proteins were pulled down by clicking onto azide-functionalized magnetic beads. Note that a small part of the lysate was saved to perform click chemistry with a TAMRA-azide for target visualization after SDS-PAGE separation and fluorescent gel scanning (Figure 5F). This approach unravelled the presence of several fluorescently labelled bands (probe labelling input, Figure 5G), indicating that KIRA6 AfBP could bind to different proteins. Moreover, the pull down was effective as it depleted the KIRA6 AfBP-linked proteins from the supernatant (Figure 5G). Competition experiments showed that the intensity of all fluorescent bands decreased when the binding of KIRA6 AfBP was outcompeted by the presence of KIRA6 in co-incubation experiments (Figure 5G). This demonstrated that proteins targeted by KIRA6 AfBP were effectively also *bona fide* KIRA6 targets. Next to identify these proteins, the KIRA6 AfBP pull downs were subjected to on-bead tryptic digestion followed by LC-MS/MS (see Experimental procedures). Table 1 lists the top 15 proteins (based on total spectrum counts) found to interact with KIRA6 AfBP.

Together, these results demonstrate that KIRA6 curtails the cancer cell inflammatory responses elicited by immunogenic treatments independently of IRE1α.

### HSP60 contributes to the activation of NF-κB signaling and CXCL8 production in response to immunogenic treatments

Having identified putative KIRA6 off-targets, we then wished to explore their possible relevance in the pro-inflammatory pathway targeted by KIRA6. We then focused our attention on the molecular chaperones HSP60 and HSP90 for different reasons; i) with the exception of actin, both molecular chaperones were the top-listed ATP-binding proteins identified by LC-MS/MS (Table 1), ii) HSP90 is a known regulator of NF-κB-driven inflammatory signaling (Thangjam et al. 2016; Qing et al. 2007; Gopalakrishnan, Matta, and Chaudhary 2013); and iii) the cytosolic fraction of HSP60 has been recently shown to promote CXCL8 production by cancer cells by positively regulating NF-κB (Chun et al. 2010).

We first assessed whether cellular stress evoked by anticancer treatments induced an upregulation of these chaperones. However, CDDP, MTX or Hyp-PDT did not elevate the overall expression of HSP60 in A375 cells (Figure 6 – figure supplement 1A) while CDDP and MTX, but not Hyp-PDT, reduced significantly the level of HSP90 (Figure 6 – figure supplement 1B). To explore their possible role, we then assessed whether silencing HSP60 or HSP90 (70-80 % knockdown efficacy, Figure 6C, Figure 6 – figure supplement 1C) could reproduce (at least in part) the effects of KIRA6 on the inhibition of CXCL8 production.

**Fig. 6.**
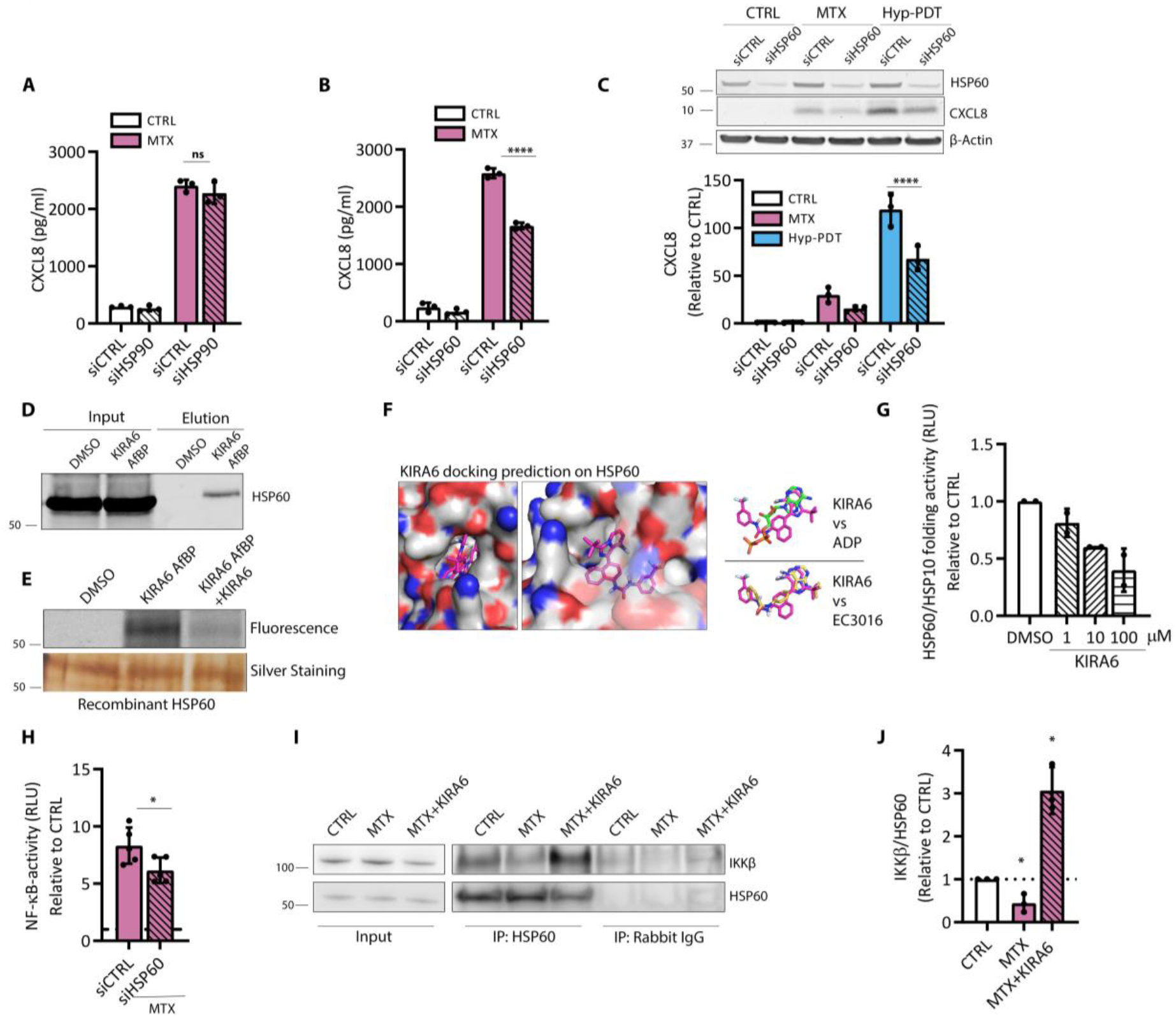
HSP60 is a KIRA6 target modulating CXCL8 production. **(A)** CXCL8 secretion was measured by ELISA in conditioned medium from A375 with siRNA mediated HSP90 knock-down (siHSP90) or **(B)** HSP60 knock-down (siHSP60) with respect to scramble siRNA (siCTRL) 24 hr after treatment with MTX. **(C)** Representative Western Blot and quantification of the impact of siHSP60 with respect to siCTRL on intracellular CXCL8 accumulation 4 hr after treatment with MTX and Hyp-PDT. Data is expressed as fold change over control incubated with siCTRL. β-actin was used as loading control. **(D** Representative Western Blot of streptavidin-mediated pull down of HSP60 with KIRA6 AfBP (10 μM) conjugated with biotin. **(E)** Representative gel of recombinant HSP60 labelling with KIRA6 AfBP (10 μM) and with co-incubation KIRA6 AfBP (10 μM) and KIRA6 (100 μM). **(F)** KIRA6 docking prediction on HSP60 (PDB: 4PJ1) using Autodock Vina. This pose is compared with that obtained from docking of EC3016 (inhibitor of GroEL, prokaryotic ortholog of HSP60) and ADP into the same structure. **(G)** In vitro refolding activity of the HSP60/HSP10 chaperone complex after 1h of incubation with heat-mediated unfolded substrate proteins in control condition or in the presence of KIRA6 at the indicated concentration; n=2 biological replicates/concentration. Data is expressed as fold change compared to control. **(H)** NF-κB activity measured by luciferase assay in A375 cells stably expressing the reporter 4 hr after treatment with MTX upon transfection with siCTRL or siHSP60. Data is expressed as fold change compared to untreated cells transfected with the correspondent siRNA, whose reference value is indicated with a dotted line. **(I-J)** Co-immunoprecipitation of IKKβ with HSP60 from the cytosolic fraction of A375 in basal condition or 2 hr after treatment with MTX or MTX coincubated with KIRA6 (10 μM). In all graphs values are presented as mean ± SD. Data is analyzed by Two-way ANOVA in all the graphs except for Student’s t-test in (H) and One-way ANOVA in (J) *p<0.05, ****p<0.0001, ns=not significant.

While the knockdown of HSP90 had no major effects (Figure 6A, Figure 6 – figure supplement 1C) HSP60 silencing reduced CXCL8 secretion after MTX (Figure 6B) and its intracellular accumulation after MTX and Hyp-PDT (Figure 6C). HSP60 depletion however did not alter the fraction of dying cells (Figure 6 – figure supplement 1D). While these results do not completely rule out the contribution of HSP90, they suggest a causal link between the anti-inflammatory action of KIRA6 and HSP60.

To further support this connection, we then evaluated whether KIRA6 AfBP was capable to covalently link this chaperone. In line with this, KIRA6 AfBP was able to pull down HSP60 from the lysate of A375 cells (Figure 6D). Furthermore, KIRA6 AfBP binding to recombinant HSP60, was competitively inhibited by the presence of KIRA6 (Figure 6E). Since KIRA6 is an ATP-competitive inhibitor, we speculated that it should bind to the nucleotide-binding site of HSP60, a notion also supported by the structural similarity between KIRA6 and two reported ATP-competitive inhibitors of HSP60 (Alagramam et al. 2016) and GroEL (Chapman et al. 2008) (prokaryotic ortholog of HSP60). In order to get an insight into KIRA6’s binding mode, we performed a docking simulation using Autodock Vina (Trott and Olson 2010) against human HSP60 (PDB: 4PJ1 (Nisemblat et al. 2015)). While the aromatic amine of the imidazopyrazine ring in KIRA6 faces the opposite direction compared with the amine of the adenine ring (ATP molecule crystallized with HSP60), we observed a good overlap between the two rings as well as good alignment with two proton acceptors (nitrogen atoms) (Figure 6F). The docking results suggest that the naphthalene moiety partially occupies the same pocket as the phosphate groups of ATP, whereas the trifluoromethyl-substituted ring occupies the adjacent pocket. This pose was also in excellent alignment with that obtained from docking EC3016 (GroEL inhibitor) into the same structure. HSP60-mediated protein folding requires ATP binding and hydrolysis (Ishida et al. 2018). To explore the biochemical effects of the docking results, we then tested the *in vitro* folding ability of recombinant HSP60 (in complex with the co-chaperone HSP10) either in the absence or the presence of different concentrations of KIRA6. HSP60 folding ability was significantly reduced by the presence of KIRA6 (Figure 6 – figure supplement 1E), in a concentration dependent manner (Figure 6G) further suggesting that the HSP60-KIRA6 complex exhibits impaired ATPase and folding ability.

We then explored the mechanism by which KIRA6 could inhibit CXCL8 production elicited by immunogenic therapy, by targeting HSP60.

HSP60 is an abundant mitochondrial chaperonin with recently reported extramitochondrial activities (Huang and Yeh 2019). We first explored whether HSP60 was relocated intracellularly after cellular stress induced by our treatments. Immunofluorescence analysis (Figure 6 – figure supplement 1F) and immunoblot (Figure 6 – figure supplement 1G) of mitochondrial and cytosolic fractions did not reveal a detectable intracellular redistribution of HSP60 after CDDP, MTX or Hyp-PDT treatments. We also failed to appreciate any significant difference in the upregulation of transcripts related to mitochondrial unfolded protein response (mtUPR) (Haynes and Ron 2010) (Figure 6 – figure supplement 1H). Altogether these results did not suggest a contribution of the mitochondrial pool of HSP60 in the pathway leading to CXCL8 production after immunogenic treatments.

HSP60 has been recently shown to regulate TNF-α-mediated activation of the NF-κB pathway via direct interaction with the IKK complex in the cytosol (Chun et al. 2010). To explore the possible inhibitory effect of KIRA6 on the HSP60-IKK complex, we then focused on the NF-κB pro-inflammatory pathway induced MTX. In line with this, HSP60 silencing partially, albeit significantly, reduced NF-κB activation in response to MTX (Figure 6H). To confirm the presence of a HSP60-IKK complex, we then performed co-immunoprecipitation of endogenous HSP60 in association with IKKα/β from the cytosolic fractions of untreated A375 cells. This co-IP analysis revealed that in unstressed cells a fraction of HSP60 was found in a complex with IKKβ (Figure 6I). Remarkably, MTX treatment (2 hr) attenuated the binding of HSP60 to IKKβ (Figure 6I, J) while the co-treatment of MTX with KIRA6 enforced the HSP60-IKKβ complex interaction (Figure 6I, J). Since MTX leads to a fast NF-κB activation, which is reduced by silencing HSP60, together these data suggest that HSP60 may initially facilitate IKK activation (probably through its folding activity), while being dispensable for its full activity. Alternatively, a fraction of the cytosolic HSP60-IKK complex could translocate to the nucleus (Chun et al. 2010; Meng, Li, and Xiao 2018). While the mechanistic aspects of this interaction requires further explorations, these findings suggest that KIRA6 by interfering with the folding activity of HSP60 locks the cytosolic HSP60-IKKβ complex in an inactive conformation and/or prevents IKKβ to dock to other partner proteins (Polley et al. 2016), thus compromising NF-κB signaling.

Altogether, these data suggest that inhibition of HSP60 by KIRA6 dampens NF-κB driven CXCL8 production and the inflammatory responses elicited by immunogenic treatments. An overall representation of the pathway is represented in Figure 7.

**Figure 7.**
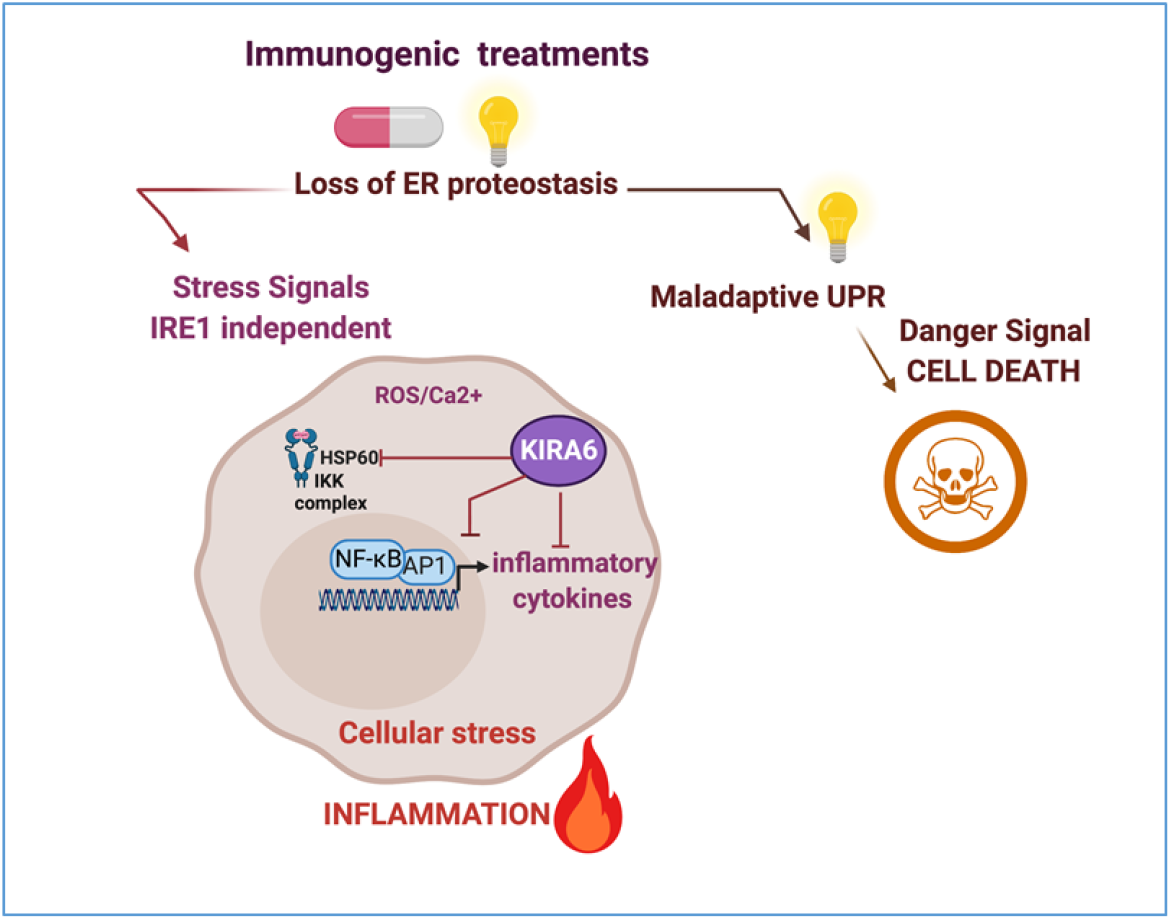
KIRA6 curtails the inflammatory traits of immunogenic treatments. Chemotherapy with mitoxanthrone or Hypericin-mediated photodynamic therapy (Hyp-PDT) (illustrated by the light as essential triggering factor) induce loss of ER homeostasis, which leads in case of the most pronounced ER stress inducer Hyp-PDT to maladaptive UPR and immunogenic cell death. Both immunogenic treatments however, also elicit an early stress response, which is independent of the UPR/ISR and caspase mediated cell death. This pre-mortem stress response leads to the production of a common subset of pro-inflammatory chemokines through ROS and Ca2+ mediated activation of the NF-κB and AP-1 transcritional program. KIRA6, an inhibitor of the IRE1α kinase activity, blunts the inflammatory output of immunogenic therapies, in a IRE1α independent manner. One of the off-target effectors of KIRA6 is the cytosolic HSP60 which is required for the full activation of NF-κB and CXCL8 production by immunogenic treatments.

## DISCUSSION

In this study, by performing RNAseq analysis and signaling pathway validation we portray an inflammatory trait shared by stressed/dying cancer cells responding to immunogenic treatments. This transcriptional signature involves NF-κB/AP-1 driven induction of a subset of pro-inflammatory chemokines, with CXCL8 emerging as a predominant entity. We show that this transcriptionally regulated pro-inflammatory pathway segregates from the activation of the UPR induced by these anticancer treatments and proceeds unrelated to caspase-induced cell death. Strikingly we show that the IRE1α inhibitor KIRA6 abolishes the inflammatory output associated to immunogenic treatments through unprecedented IRE1α independent mechanisms, which involves HSP60 modulation of NF-κB driven CXCL8 production (Figure 7).

The causal link between the UPR pathway and sterile inflammation has been elucidated in several studies (Schmitz et al. 2018; Hotamisligil 2010; Zhang and Kaufman 2008; Keestra-Gounder et al. 2016) using both genetic and pharmacological approaches, which provided independent support for the role of the PERK and IRE1α pathways in NF-κB activation. Moreover, recent studies proposed that canonical ER stress agents, such as thapsigargin or tunycamicin, but also chemotherapies such as taxanes, stimulate the PERK/ATF4/CHOP-dependent upregulation of DR5 (TRAIL-R2) (Lu et al. 2014; Iurlaro et al. 2017; Martín-Pérez et al. 2014; Glab et al. 2017; Lam et al. 2020). DR5 upregulation then causes ligand independent FADD/caspase-8/RIPK1-driven NF-κB-activation and pro-inflammatory signaling (Sullivan et al. 2020). In this model, DR5 operates as a sensor of misfolded proteins, which act as ligands for DR5 in the ER-Golgi intermediate compartment (Lam et al. 2020), thereby launching inflammatory responses aimed at restoring homeostasis (Sullivan et al. 2020).

Our study however shows that this is not a general ‘modus operandi’ of all agents eliciting ER stress. Here we studied prototypes of immunogenic treatments known to stimulate in the stressed/dying cancer cells, loss of ER proteostasis associated to the concomitant extracellular redistribution of pre-existing endogenous molecules, or DAMPs, and the secretion of immunostimulatory factors, like type I interferon, which together ultimately license antitumor immunity (Kroemer, Galluzzi, and Zitvogel 2016; Garg and Agostinis 2017). We found that MTX and prevalently Hyp-PDT (which induces a more persistent pro-apoptotic PERK signal (Garg et al. 2012; Verfaillie et al. 2012)) upregulate DR5 and induce caspase signaling, but both signals are dispensable for the transcriptional upregulation of CXCL8. A differential involvement of the CHOP-DR5 pathway could be inherent in the apical triggers causing loss of ER homeostasis and consequent UPR or ISR. The immunogenic treatments explored in our studies, promote a rapid NF-κB and AP-1 activation as an early (pre-mortem) stress responses stimulated by Ca^2+^ and ROS-induced signals, which largely precede apoptotic cell death. In fact, in spite of being a type II topoisomerase inhibitor and supposedly inhibiting DNA and RNA synthesis, MTX elicits various extra-nuclear activities, including ER-Ca^2+^ depletion and ROS, which drive eIF2α-phosphorylation and immunogenic cell death (Garg et al. 2012; Panaretakis et al. 2009; Martins et al. 2011). The common generation of ROS by these treatments may impair assembly of the DR5-caspase8 complex at the ER-Golgi compartment or induce redox modification of the misfolded proteins, thereby diminishing their ability to serve as DR5 ligands. While this hypothesis requires further studies, PDT using hypericin or redaporfirin as photosensitizers results in ROS-induced perturbations of the ER-Golgi compartment (Gomes-da-Silva et al. 2018; Garg et al. 2012) causing a general inhibition of protein and cytokine secretion as observed here and in our previous study (Dudek-Peric et al. 2015). Whether and how the intracellular accumulation of cytokines by PDT influences their pro-inflammatory and immunomodulatory output when cell death ultimately ensues need further evaluation *in vivo*.

Irrespective of this unknown, our transcriptomic profiling and pathway analysis clearly show that in our settings Ca^2+^ and ROS-sensitive signals elicit pro-inflammatory responses independent of the activation of the UPR pathway. Indeed, we demonstrate that the unique ability of the IRE1α kinase inhibitor, KIRA6, to overrule the pro-inflammatory output of immunogenic treatments occurs through a mechanism that is independent of the presence of IRE1α.

KIRA6 is a small molecule inhibitor and prototype of ATP-binding compounds which disrupt IRE1α oligomerization thereby suppressing the RNase activities of IRE1α (Harnoss et al. 2019; Ghosh et al. 2014). Since its development (Ghosh et al. 2014) KIRA6 has been used in several studies as specific inhibitor of the IRE1α branch of the UPR-associated inflammatory responses (Keestra-Gounder et al. 2016) and *in vivo* to prevent inflammation-driven pathologies (Thamsen et al. 2019; Ghosh et al. 2014).

In line with its proposed anti-inflammatory role, in chemotherapy (MTX) treated cancer cells of human and murine origin, KIRA6 potently inhibited NF-κB driven transcription, secretion of several cytokines/chemokines and CXCL8-mediated neutrophil chemotaxis. In this study we used a KIRA6-clickable photoaffinity probe which we recently generated to identify functionally relevant off-targets of this compound (Korovesis et al. 2020). Using this approach, we revealed that KIRA6 is able to inhibit a signaling circuit involving the cytosolic pool of HSP60 in complex with IKKβ. Functional, biochemical and co-IP assays, further revealed that KIRA6 by targeting HSP60 reduces its folding activity and in MTX treated cancer cells locks the cytosolic HSP60-IKKβ complex likely in a less active conformation, with a reduced ability to activate NF-κB mediated pro-inflammatory responses.

HSPs respond to loss of proteostasis by exerting various intracellular cytoprotective functions and by regulating signaling pathways that play critical roles in inflammation and immune responses, therefore the term chaperokines (Sevin et al. 2015). Key regulators of inflammatory and immune responses are clients of cytosolic HSP90, and HSP90 inhibitors such as geldanamycin and radicicol that bind to its ATP pocket, are promising anticancer therapeutics (Sevin et al. 2015). In contrast, a role for the cytosolic pool of HSP60 in the NF-κB pathway has only recently emerged (Chun et al. 2010). While further investigations are required to fully appreciate the mechanistic underpinnings of the inhibitory effects of KIRA6, our study highlights a preferential role for HSP60 in the pro-inflammatory pathway initiated by immunogenic treatments, thereby revealing a previously unexplored intracellular role for HSP60 elicited by these anticancer therapies.

In conclusion, we show that immunogenic treatments share a common transcriptional signature, involving a pro-inflammatory early stress response operating in parallel to the UPR and which can be overruled by KIRA6, independently of IRE1α. This unexpected finding is relevant, considering that small molecule inhibitors of the IRE1α pathway have been proposed as potential anticancer therapeutics (Hetz, Chevet, and Harding 2013), due to their ability to inhibit inflammation and the recruitment of myeloid immune cells within the tumor. Beyond anticancer treatments, our study raises caution about the use of KIRA6 to assess the role of IRE1α in inflammatory pathologies.

## Acknowledgments

We would like to thank Maarten Ganne for excellent technical assistance. P.A. is supported by grants from the Flemish Research Foundation (FWO-Vlaanderen; G076617N, G049817N, G070115N), the EOS consortium (30837538, with A.D.G.) and Stichting tegen Kanker (FAF-F/2018/1252) and KU Leuven (C16/15/073). N.R. was funded by European Union’s Horizon 2020 research and innovation programme under the Marie Sklodowska-Curie grant agreement no. 642295. M.V.P. is the recipient of an FWO Doctoral Fellowship from the Flemish Research Foundation (FWO-Vlaanderen, 1186019N), Belgium. F.F. is supported by the Austrian Science Fund (FWF) (project No. T 974-B30).

Images were recorded on a Zeiss LSM 780 – SP Mai Tai HP DS (Cell and Tissue Imaging Cluster, CIC), supported by Hercules A KUL/11/37 and FWO G.0929.15 to Pieter Vanden Berghe, University of Leuven.

## Materials and Methods

### Chemical inhibitors

Brefeldin A, SCH772984, 4μ8C and KIRA6 were purchased by Selleckchem. BAY 11-7082, GSK2606414, STF-083010 were purchased by Caymanchem). N-Acetyl-L-cysteine and L-Histidine were supplied by Sigma-Aldrich, thapsigargin by Enzo Life Sciences, BAPTA-AM by ThermoFisher Scientific. Bortezomib was obtained directly from the Pharmacy Department of the Leuven University Hospital, Leuven Belgium,.

### Cell culture and treatments

A375 were cultured in DMEM (Sigma-Aldrich) supplemented with 2mM glutamine (Sigma-Aldrich) and 10% fetal bovine serum (PAN-Biotech); CT26, MEF, HeLa, HCT116, THP1 and HEK293T were cultured in DMEM supplemented with 1 mM glutamine and 10% FBS. All cell lines were cultured at 37°C under 5% CO2. To induce cell death A375, HeLa, MEF and HCT116 cells were incubated with 1 μM mitoxantrone (MTX, Sigma-Aldrich) or 50 μM Cisplatin (CDDP, Sigma-Aldrich). CT26 were incubated with 5 μM MTX and 50μM CDDP. For hypericin-based photodynamic therapy (Hyp-PDT) conditions, A375 cells were incubated for 16 hr with 150 nM hypericin (Enzo Life Sciences) in full medium followed by removal of hypericin, irradiation with fluence=2.70 J/cm2, and cultured for indicated times. Whenever chemical inhibitors were used, they were preincubated for 1h before addition of cell death inducers and maintained in the medium during cell death induction.

### Cell death assay

For cell death kinetics, cells were treated with cell death inducers in presence or absence of 50 μM z-Vad-FMK (Bachem) in medium containing 1 μM sytox green (Thermofisher Scientific). At the indicated time post-treatment, fluorescence emission at λ=530nm was measured at flexstation 3 (Molecular Devices). Cells were then lysed with Cell lysis buffer (Bioke) for 10 minutes at room temperature (RT) and fluorescence relative to 100% cell death was measured and used as normalization parameter. For other cell death assays, cells were collected 24 hr after treatment with the cell death inducers with TrypLE Express (Life Technologies) and resuspended in PBS containing 0.5% bovine serum albumin (BSA, Sigma-Aldrich) and 1 μM sytox green and analyzed by flow cytometry on Attune (Thermofisher Scientific).

### Ecto-Calreticulin detection

After treatment, cells were collected with TrypLE Express (Life Technologies), washed with PBS and with Flow cytometry (FC) buffer (3% BSA in PBS), incubated for 30 minutes at 4°C with fluorophore-conjugated primary antibody (EPR3924, AbCAM), washed with FC buffer and resuspended in FC buffer including viability die and analyzed on Attune (Thermofisher Scientific). The permeabilized cells were excluded from the analysis due to intracellular staining.

### ATP assay

A375 cells were treated as indicated in 2% FBS medium. Extracellular ATP was measured in the conditioned with ATP Bioluminescent assay kit (Sigma-Aldrich) following manufacturer’s instructions. Bioluminescence was assessed by optical top reading via FlexStation 3 microplate reader (Molecular Devices).

### Immunofluorescence

Cells were plated on coverslips precoated with 0.1% gelatin, treated for the indicated time and fixed in 4% PFA. Cells were permeabilized 10 minutes with 0.1% Triton in PBS, then blocked for 1 hr in blocking buffer (5% FBS, 1% BSA) for 1 hr. Primary antibody was added in blocking solution and incubated overnight at 4°C. After washing with TBS-T buffer (50 mM Tris, 150 mM NaCl, 0.1% Tween-20), coverslips were incubated with secondary antibody for 2h at RT. Antibodies used are listed in Supplementary Table 1. Nuclei were counterstained with DAPI in PBS (1 μg/ul, 1:1000) and mounted on slides with Prolong Gold. Pictures were taken at Zeiss LSM 780 confocal microscope. For the representative image, the central stack was selected using ImageJ software.

### Western Blot

Whole cell lysates were loaded on 4-12% Bis-Tris gel and separated by SDS-PAGE on the Criterion system (Bio-Rad Laboratories), and electrophoretically transferred to nitrocellulose membrane. The blots were blocked for 1 hr at RT with in TBS-T buffer containing 5% non-fat dry milk, and incubated with the indicated primary antibodies overnight at 4°C in TBS-T containing 5% BSA. After washing with TBS-T, membranes were incubated with secondary antibodies conjugated with infrared fluorophores for 2 hr at RT. The membranes were visualized with Typhoon biomolecular imager (Amersham). Alternatively, horseradish peroxidase secondary antibodies were used and the membranes were incubated with enhanced luminol-based chemiluminescent substrate (Amersham) and visualized with Amersham Imager 600. All antibodies used are listed in Supplementary Table 1.

### RNA extraction and qPCR

After treatment for the indicated time, total RNA was extracted from cells using TriSure buffer (Bioline) followed by phenol/chloroform extraction and 1 μg of RNA was reverse transcribed with Quantitect RT kit (Qiagen) following manufacturer’s instructions. Primers for real-time PCR were designed with the Primer3 web tool (Supplementary Table 2). The housekeeping 18S ribosomal RNA was used to normalize the expression levels. Quantitative PCR (qPCR) was performed with ABI 7500 Fast Real-Time PCR System using ORA™ qPCR Green Master Mix (highQu).

### XBP1 splicing assay

The following primers were used: unspliced XBP1 (5’-CAGCACTCAGACTACGTGCA-3’, sense), spliced XBP1 (5’-CTGAGTCCGAATCAGGTGCAG-3’, sense), unspliced and spliced XBP1 (5’- ATCCATGGGGAGATGTTCTGG-3’, antisense). The RT-PCR analyses were performed according to the following conditions: denaturation at 95°C for 10 minutes, followed by 35-cycles of denaturation at 95°C for 10 seconds, annealing at 5 °C for 30 seconds, extension at 7 °C for 1 minute and final extension at 72 °C for 7 minutes. Amplicons were resolved in 2% agarose gels.

### ELISA

Eight or 24 hr after treatment, conditioned medium was collected and the levels of secreted human CXCL8 and murine CXCL1 were detected with DuoSet ELISA (R&D systems) following the manufacturer’s instructions. Absorbance values were measures on Flexstation 3 at λ=530nm and absorbance background values at λ=540nm were subtracted.

Multiplexed ELISA was performed on a Luminex FLEXMAP 3D^®^ platform (Luminex, Austin, TX), using a custom-developed chemokine 6-plex panel (ProtATonce, Athens, Greece). Custom antibody-coupled beads were technically validated as described before (Poussin et al. 2014).

### SiRNA transfection

Cells were transfected by adding 1 ml serum-free culture media with Trans-IT X2 transfection reagent (MirusBio) and targeting (On-Target Smartpool siRNA) or scrambled (siCTRL) siRNA (Dharmacon, Thermo Fisher Scientific) twice on two consecutive days. Experiments were performed 48 hr after the second transfection.

### NF-κB activity

Plasmid pHAGE NFkB-TA-LUC-UBC-GFP-W containing the luciferase gene under the minimal NF-κB promoter was a gift from Darrell Kotton (Addgene plasmid #49343; http://n2t.net/addgene:49343; RRID:Addgene_49343). To generate lentiviral particles, HEK 293T cells were transfected with pHAGE plasmid in the presence of plasmid encoding VSV-G (pMD2-VSV-G, Tronolab) and packaging proteins (pCMVdR8.9, Tronolab). Twenty-four hr after transfection transfecting medium was substituted with fresh medium and VSV-G pseudotyped virus was collected 48 hr after transfection and added to the exponentially growing A375 cell cultures in the presence of hexadimethrine bromide (Sigma-Aldrich). Cells were expanded and GFP positive cells were sorted with Influx. Four hr post-treatment, NF-κB activity was measured with luciferase assay kit (BioAssay System) following manufacturer’s protocol at flexstation3.

### Chemotaxis

Human neutrophils were obtained from fresh human peripheral blood from healthy volunteers and were isolated using a Percoll (Sigma-Aldrich) gradient.

Conditioned supernatants were generated from treated A375 cells for 24 hr in DMEM without FBS. Conditioned supernatants were then deprived of chemical inhibitors by using Amicon^®^ Centrifugal Filter Unit and washed with two volumes of PBS and resuspended to 10x the initial concentration. Chemotaxis assays were performed with Transwell^®^ polycarbonate membrane cell culture inserts (Corning). 100 μL of 10-times concentrated supernatant diluted in 400 μL of DMEM were added to the bottom well of a chemotaxis chamber and 5×10^4^ THP-1 cells or primary human peripheral blood neutrophils were added. For rescue experiments, CXCL8 and CXCL8 neutralizing antibody were added in the bottom well at a concentration of 10 ng/ml 0.5 μg/ml, respectively. Total cells entering the bottom chamber were counted after 2 hr and representative pictures were taken at Olympus IX73 inverted microscope (Olympus Life Sciences).

### IRE1 knock-out generation

A375 IRE1 knock-out and scramble control were generated with CRISPR-Cas9 double nickase system (Santa Cruz Biotechnology) according to manufacturer’s protocol.

### KIRA6 AfBP generation and target labeling

KIRA6 AfBP was generated as previously reported (Korovesis et al. 2020).

A375 whole cell lysates were normalized to a concentration of 1 mg/mL in a volume of 30 μL. Samples were then treated with KIRA6 AfBP (10 μM) or DMSO, mixed by vortexing and immediately irradiated for 6 minutes at RT. For competition experiments, samples were co-treated with probe (10 μM) and KIRA6 (100 μM). For HSP60 labelling, 160 ng of recombinant human HSP60 were used.

After irradiation, probes were clicked onto TAMRA-azide (Carl Roth) using the following conditions: 25 μM of tag-azide, 50 μM of THPTA (Sigma-Aldrich Aldrich), 1 mM of CuSO_4_ (freshly prepared) and 1 mM of sodium ascorbate (freshly prepared). Click reaction was incubated for 1 hr at RT and the reaction was quenched by addition of 10 μL of 4x SDS loading buffer. Samples were resolved by 10% SDS-PAGE. Following visualization, gels were stained with coomassie using ROTI^®^ Blue (Carl Roth).

### Mass spectrometry and data analysis

Live A375 cells were incubated with KIRA6 AfBP (10 μM) for 1 hr and irradiated for 6 minutes at RT. Proteins were then extracted and probes were clicked onto azide-functionalized magnetic beads (Jena Bioscience) as described above. Probe-labelled proteins were enriched with magnetic isolation and extensively washed. To perform disulfide bonds reduction and cysteine alkylation, the beads were resuspended in denaturation buffer (7 M urea, 20 mM HEPES) and DTT (1 mM) was added for 45 minutes at room temperature. Then iodoacetamide (4 mM) was added and incubated 45 minutes at room temperature. Finally, DTT (5 mM) was added for 45 minutes at room temperature to quench the remaining iodoacetamide. On-bead trypsin digestion was executed overnight at 37 °C in the presence of 0.6 μg of trypsin, 200 mM ammonium bicarbonate, 2.5 % acetonitrile and 0.005 % ProteaseMax. The resulting peptide mixture was subjected to C18 Zip Tip clean-up (Millipore) before being analyzed by high-resolution LC-MS/MS using an Ultimate 3000 Nano Ultra High Pressure Chromatography (UPLC) system interfaced with an orbitrap Elite mass spectrometer via an EASY-spray (C-18, 15 cm) column (Thermo Fisher Scientific). Peptides were identified by MASCOT (Matrix Science) using the Homo sapiens database (173330 entries), adopting the following MASCOT search parameters: trypsin, two missed cleavages allowed, oxidation (M) was specified as variable modification, carbamidomethylation of cysteine was specified as fixed modification. Mascot was searched with a fragment ion mass tolerance of 0,50 Da and a parent ion mass tolerance of 10 ppm. Scaffold software was used to validate MS/MS based peptide and protein identifications, being accepted if they could be established by a probability greater than 95% and 99%, respectively. The presence of at least two unique identified peptides per protein was required.

### Off-target validation by pull down

After coincubation with A375 protein lysates and irradiation, probes were clicked onto TAMRA-azide-PEG-biotin as described above. The excess reagents from the samples were then removed by acetone precipitation. Following resuspension of the pellets to a final volume of 100 μL, half of the sample was kept as the input control. The remaining 50 μL were incubated with 20 μL of pre-washed streptavidin beads (ThermoFisher) for 1 hr with mixing at RT. The supernatant was removed and beads were sequentially washed with 0.33% SDS in PBS, 1 M NaCl and PBS. Bound proteins were eluted with sample buffer (62.5 μM Tris-HCl, 10% glycerol, 2% SDS, 1x protease inhibitor, 1x phosphatase inhibitor) and resolved by Western Blot.

### RNAseq and bioinformatics analyses

Raw RNAseq FASTQ files were preprocessed to remove technical artifacts, performing quality trimming to trim low-quality ends (< Q20) and remove trimmed reads shorter than 35bp using FastX 0.0.14 (Gordon and Hannon 2010), adapter trimming (considering at least 10bp overlap and 90% match) with cutadapt 1.7.1 (Marcel Martin 2011), and quality filtering using FastX 0.0.14 and ShortRead 1.24.0 to remove polyA-reads, ambiguous reads containing N’s, low quality reads (with more than 50% of the bases < Q25) and artifact reads (with all but 3 bases in the read equal one base type). RNAseq reads were aligned to the Homo sapiens GRCh3773 reference genome using STAR 2.4.1d (Dobin et al. 2013), with the following parameter settings: --outSAMprimaryFlag OneBestScore ‒twopassMode Basic -- alignIntronMin 50 --alignIntronMax 500000 --outSAMtype BAM SortedByCoordinate. The samtools 1.1 toolkit was used to remove reads with non-primary mappings or with a mapping quality ≤ 20 (Li et al. 2009) and for BAM/SAM file sorting and indexing. Gene counts were computed with featureCounts 1.4.6 (Yang Liao, Smyth, and Shi 2014), using the following options: -Q 0 -s 2 -t exon -g gene id. RNAseq data are deposited in the GEO repository with accession number GSE163377.

Differential expression analysis was performed with the R package edgeR (Robinson, McCarthy, and Smyth 2009), considering only protein-coding genes with at least one count-per-million (CPM) in not less than three samples. Gene counts were normalized between samples with trimmed-mean of M-values (TMM) normalization (Robinson and Oshlack 2010) and dispersions were estimated with the Cox-Reid profile-adjusted likelihood method (McCarthy, Chen, and Smyth 2012). Given the small number of replicates, the quasi-likelihood F-test was used for testing (Lun, Chen, and Smyth 2016). Genes differentially expressed in the treated cell lines with respect to the control at any time point were selected using a level of significance of 0.1 on p-values adjusted for multiple testing with the Benjamini-Hochberg approach and imposing a cut-off of one on absolute log-fold-changes. Principal component analysis was performed on the 10% most variable genes considering rlog-normalized counts computed with the DeSeq2 R package (Love, Huber, and Anders 2014). Gene Ontology (GO) was performed with the R package *TopGO* (Alexa and Rahnenfuhrer 2016) using as input the significantly upregulated genes compared to time matched untreated control separately by each treatment at each time point analyzed and using the entire gene list mapped by RNAseq as reference library. The gene ontology annotations relative to Biological Processes and Cellular Compartments were provided by the *org.Hs.eg.db* and *GO.db* annotation packages. Gene Set Enrichment Analysis (GSEA) (Subramanian et al. 2005) was perform on WebGestalt (Yuxing Liao et al. 2019) against Wikipathway Cancer database submitting as entry the output from PCA with genes weighed for their contribution to the variance of the first two principal components. Transcription factor prediction was performed with Ingenuity Pathway Analysis (Qiagen), iRegulon (Janky et al. 2014) Cytoscape plugin and TRANSFAC annotation-based GATHER (Chang and Nevins 2006) tool submitting as entry the list of genes belonging to the GO Cellular Component term “extracellular space” that were jointly upregulated by Hyp-PDT and MTX treatments.

### Cytosol isolation

Cytosol and mitochondria were separated from A375 cells 2 hr or 4 hr after treatment using the Mitochondria/Cytosol Fractionation Kit (Abcam) following the manufacturer’s instructions.

### Co-Immunoprecipitation

Cytosol was isolated as described above from A375 2 hr after treatment and 500 μg of proteins were combined with primary antibody overnight at 4°C. Protein-antibody complexes were captured by addition of Protein AG Magnetic Beads (Pierce) for 1.5 hr at RT. Protein AG Magnetic Beads with captured protein-antibody complexes were washed three times with lysis buffer. Proteins were eluted with sample buffer (62.5 μM Tris-HCl, 10% glycerol, 2% SDS, 1x protease inhibitor, 1x phosphatase inhibitor) and loaded on gel for western blot analysis.

### HSP60 refolding assay

Protein refolding efficiency of HSP60/HSP10 chaperone complex was assessed with Human HSP60/HSP10 Protein Refolding Kit (Biotechne) following manufacturer’s instructions.

---

**Supplementary Table 1.**
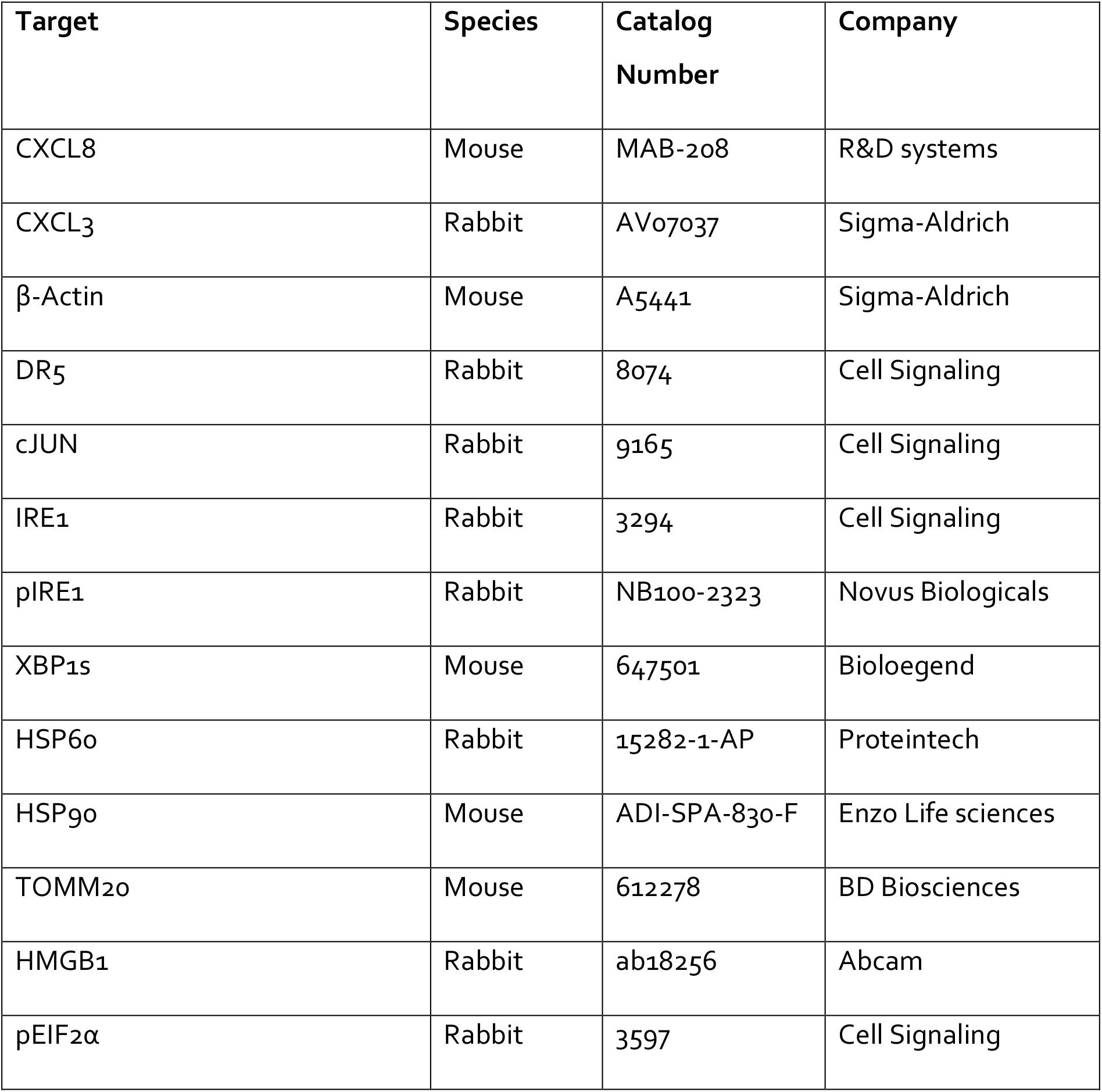
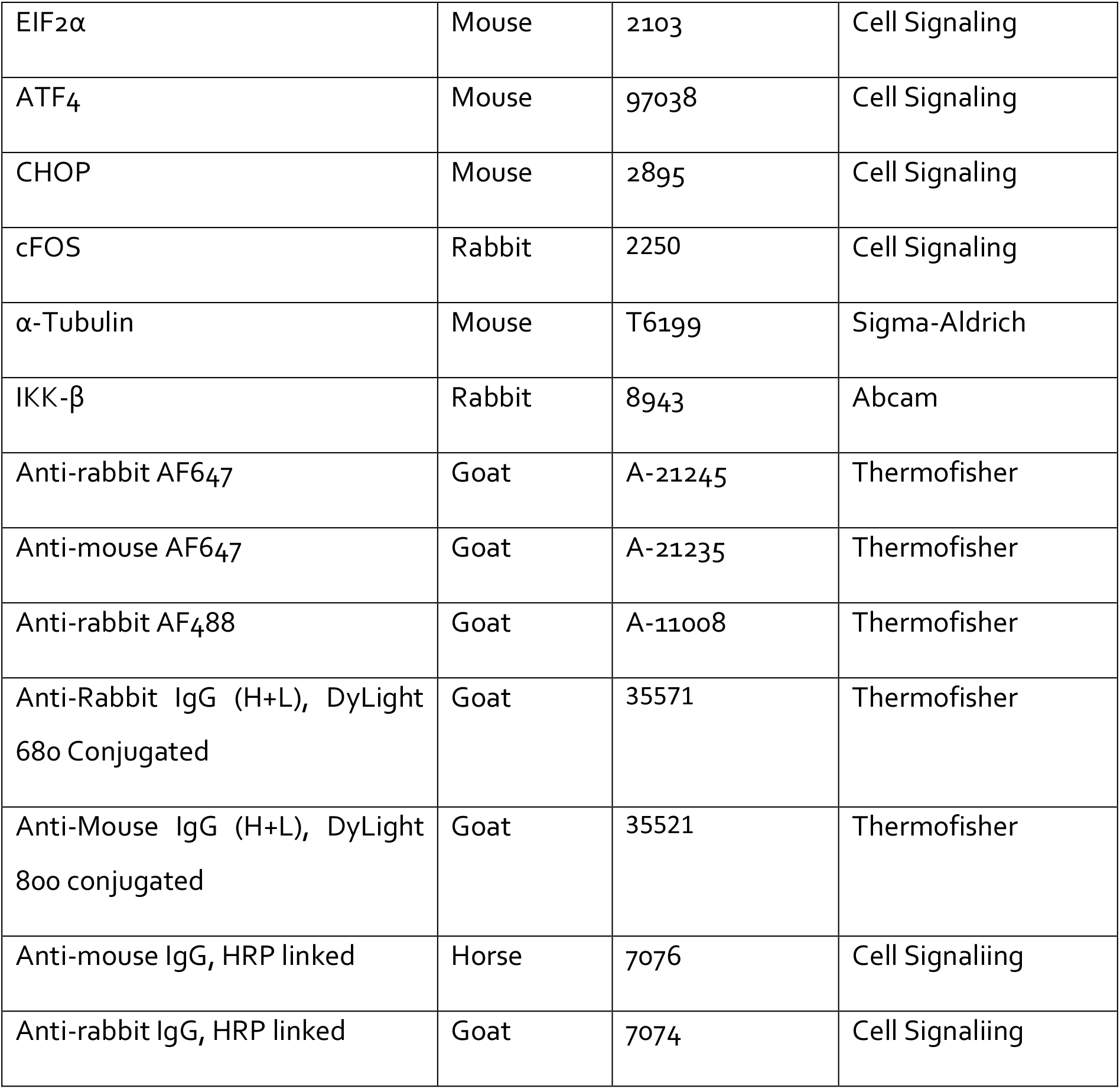
Antibodies.

| Target | Species | Catalog Number | Company |
| --- | --- | --- | --- |
| CXCL8 | Mouse | MAB-208 | R&D systems |
| CXCL3 | Rabbit | AV07037 | Sigma-Aldrich |
| β-Actin | Mouse | A5441 | Sigma-Aldrich |
| DR5 | Rabbit | 8074 | Cell Signaling |
| cJUN | Rabbit | 9165 | Cell Signaling |
| IRE1 | Rabbit | 3294 | Cell Signaling |
| pIRE1 | Rabbit | NB100-2323 | Novus Biologicals |
| XPB1s | Mouse | 647501 | Biologend |
| HSP60 | Rabbit | 15282-1-AP | Proteintech |
| HSP90 | Mouse | ADI-SPA-830-F | Enzo Life sciences |
| TOMM20 | Mouse | 612278 | BD Biosciences |
| HMGB1 | Rabbit | ab18256 | Abcam |
| pEIF2α | Rabbit | 3597 | Cell Signaling |
| EIF2α | Mouse | 2103 | Cell Signaling |
| ATF4 | Mouse | 97038 | Cell Signaling |
| CHOP | Mouse | 2895 | Cell Signaling |
| cFOS | Rabbit | 2250 | Cell Signaling |
| α-Tubulin | Mouse | T6199 | Sigma-Aldrich |
| IKK-β | Rabbit | 8943 | Abcam |
| Anti-rabbit AF647 | Goat | A-21245 | Thermofisher |
| Anti-mouse AF647 | Goat | A-21235 | Thermofisher |
| Anti-rabbit AF488 | Goat | A-11008 | Thermofisher |
| Anti-Rabbit IgG (H+L), DyLight 680 Conjugated | Goat | 35571 | Thermofisher |
| Anti-Mouse IgG (H+L), DyLight 800 conjugated | Goat | 35521 | Thermofisher |
| Anti-mouse IgG, HRP linked | Horse | 7076 | Cell Signaling |
| Anti-rabbit IgG, HRP linked | Goat | 7074 | Cell Signaling |

**Supplementary Table 2.**
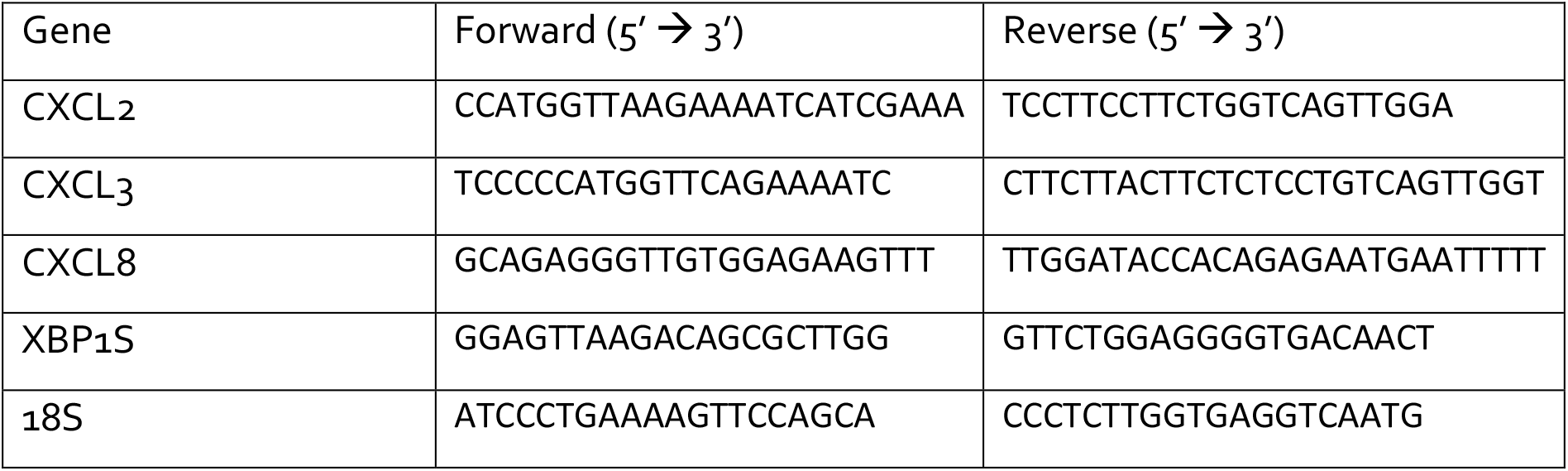
PCR Primers.

## Supplementary Figures

**Figure 1 – figure supplement 1.**
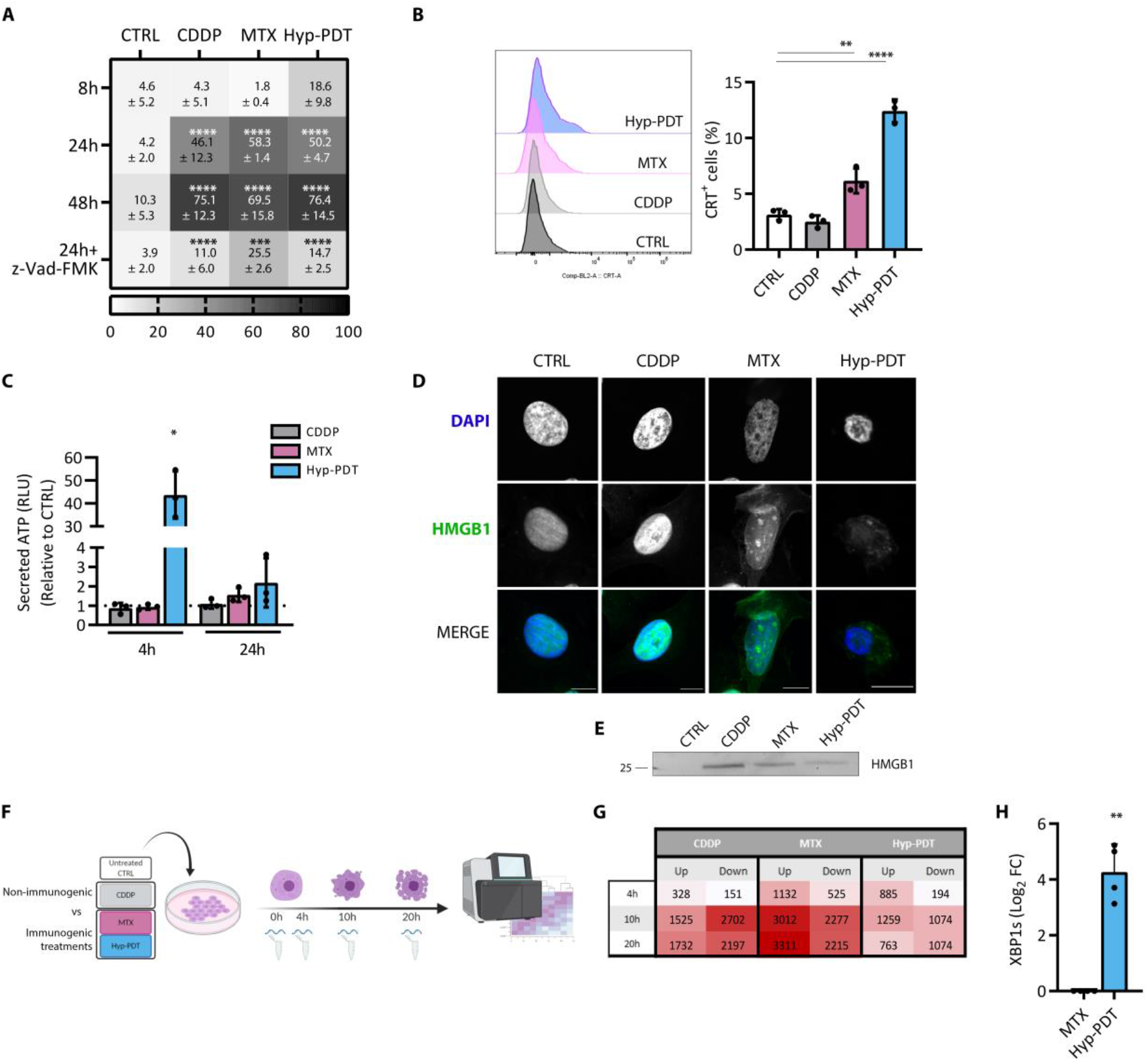
**(A)** Heatmap summarizing percentage of cell death measured at the indicated time points after treatment of A375 cells with CDDP, MTX or Hyp-PDT detected by the uptake of Sytox Green. The impact of the pan-caspase inhibitor z-Vad-FMK (50 μM) on cell death in basal condition or induced by CDDP, MTX and Hyp-PDT was assessed 24 hr after treatment. N=3 independent biological replicates **(B)** A375 cells treated with CDDP, MTX or Hyp-PDT were evaluated for the externalization of CRT in non-permeabilized cells by FACS staining 4 hr after treatment. Dead cells were excluded from the analysis. **(C)** ATP secretion in the medium was measured by luciferase-based assay at the indicated time points after treatment with CDDP, MTX and Hyp-PDT. **(D)** Representative confocal microscopy images for the intracellular redistribution of HMGB1 (Green) in A375 cells 16 hr after treatment with CDDP, MTX and Hyp-PDT. Nuclei were counterstained with DAPI (blue). Scale bar: 10 μM. **(E)** Representative Western Blot for HMGB1 secretion in the medium from A375 24 hr after treatment with CDDP, MTX and Hyp-PDT. **(F)** Schematic representation of the experimental setup utilized for the RNAseq. **(G)** Heatmap summarizing the significantly upregulated and downregulated genes identified by RNAseq at the indicated time after treatment with CDDP, MTX and Hyp-PDT. Significance was set with FDR<0.01 compared to time-matched untreated control and cut-off of 1 on log2-fold changes. **(H)** mRNA levels of cleaved XBP1 (XBP1s) were measured by qPCR in A375 cells 4 hr after treatment with MTX and Hyp-PDT. In all graphs values are presented as mean ± SD. Data is analyzed by two-way ANOVA in (A), one-way ANOVA in (B, C) and one sample t-test in (H), *p<0.05,**p<0.01, ***p<0.001, ****p<0.0001 versus time-matched control, $p<0.05, $$p<0.001 versus time matched-treatment.

**Figure 2 – figure supplement 1.**
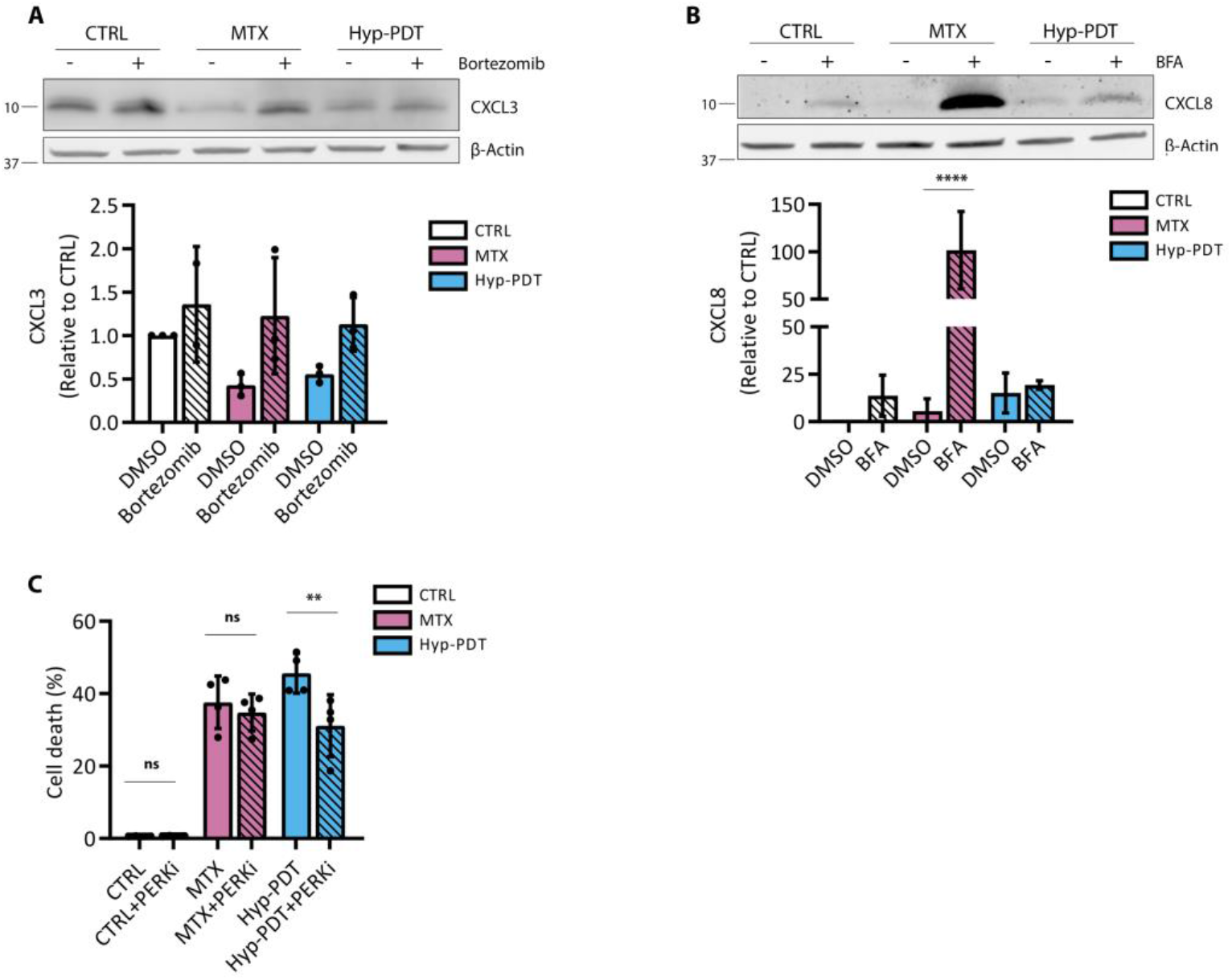
**(A)** Intracellular CXCL3 levels were assessed 8 hr after treatment with MTX and Hyp-PDT in the presence or absence of the proteasome inhibitor Bortezomib (150 nM). **(B)** Intracellular CXCL8 levels were assessed 24 hr after treatment with MTX and Hyp-PDT in the presence or absence of Brefeldin A (BFA, 50 ng/ml). **(C)** Cell death was assessed 24 hr after treatment with MTX and Hyp-PDT in the presence of absence of PERK inhibitor GSK2606414 (1 μM In Western Blots β-actin was used as loading control. In all graphs values are presented as mean ± SD of three independent experiments. Data is analyzed by Two-way ANOVA, **p<0.01, ****p<0.0001, ns=not significant.

**Figure 3 – figure supplement 1.**
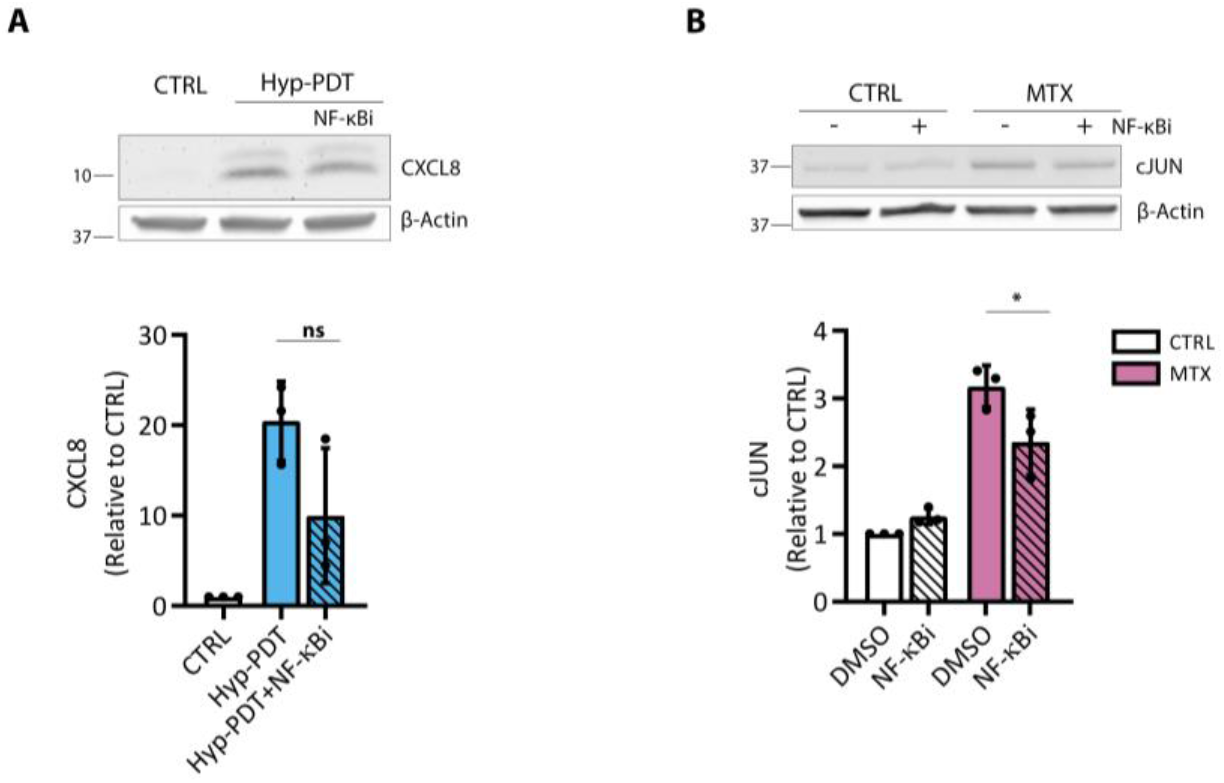
**(A)** Intracellular CXCL8 levels were assessed 4 hr after treatment with Hyp-PDT in the presence or absence of NF-κB inhibitor BAY11-7082 (10 μM). **(B)** Intracellular levels of cJUN were evaluated 4 hr after treatment with MTX in the presence or absence of NF-κB inhibitor BAY11-7082 (10 μM). In all Western Blots β-actin was used as loading control. Data is presented as mean ± SD and analyzed by Student’s t-test in (A) and Two-way ANOVA in (B), *p<0.05, ns=not significant.

**Figure 4 – figure supplement 1.**
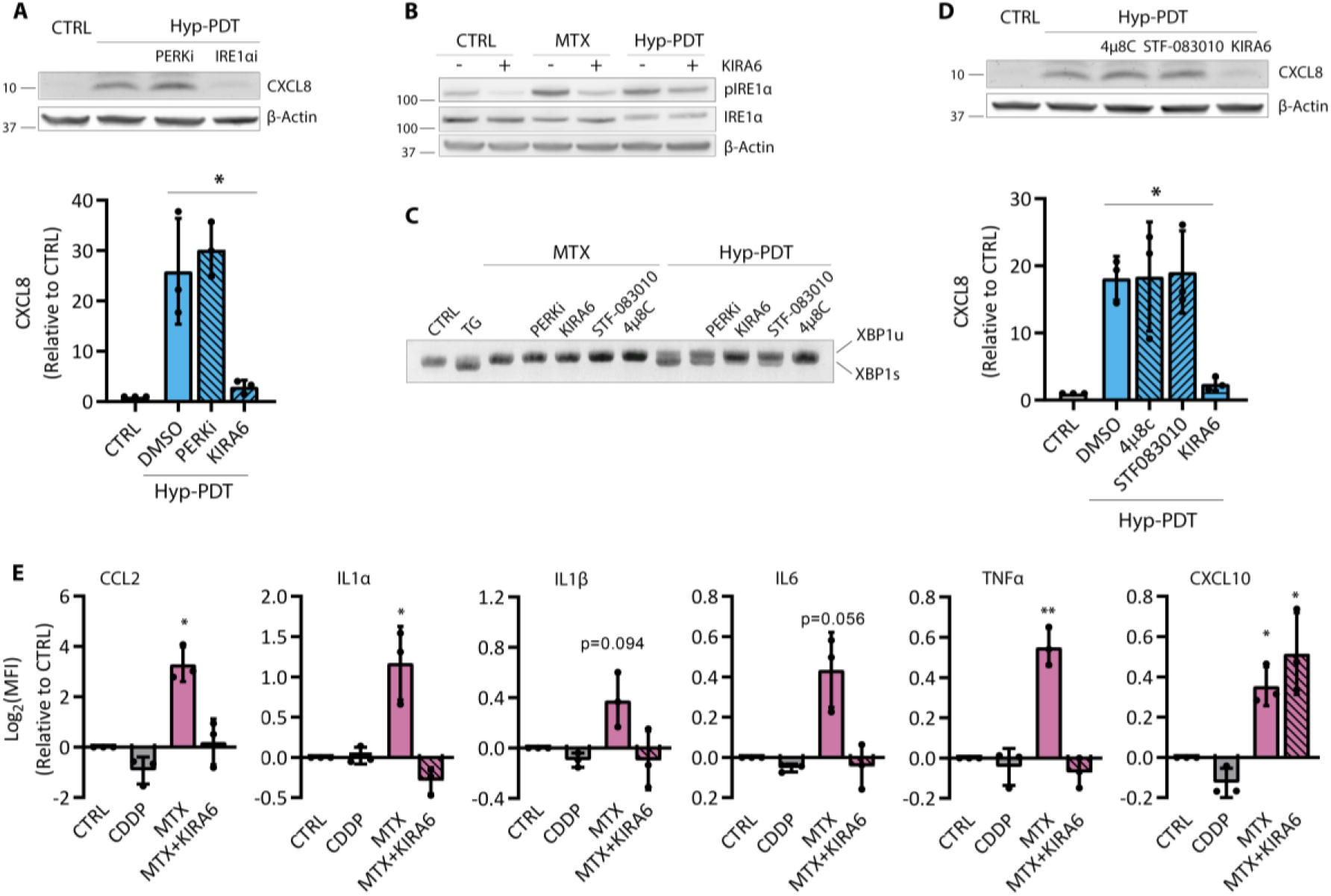
**(A)** Intracellular CXCL8 levels were assessed 4 hr after treatment with Hyp-PDT in the presence or absence of the PERK inhibitor GSK2606414 (1 μM) and the IRE1α inhibitor KIRA6 (1 μM). **(B)** Representative Western blot evaluating the efficacy of the IRE1α kinase inhibitor KIRA6 (1 μM) in inhibiting phosphorylation of IRE1α and intracellular CXCL8 accumulation 4 hr after treatment with MTX and Hyp-PDT. **(C)** XBP1 splicing assay was performed by PCR 4 hr after treatment with MTX and Hyp-PDT in the presence or absence of the PERK inhibitor GSK2606414 (1 μM) and the IRE1α inhibitors 4μ8C (100 μM), STF-083010 and KIRA6 (1 μM). The upper band refers to unspliced XBP1 (XBP1u) and the lower band to spliced XBP1 (XBP1s). Thapsigargin (TG, 2 μM) was used as positive control for XBP1 splicing. **(D)** Intracellular CXCL8 levels were assessed 4 hr after treatment with Hyp-PDT in the presence or absence of the IRE1α RNAse inhibitors 4μ8C (100 μM) and STF-083010 (50 μM) and IRE1α kinase inhibitor KIRA6 (1 μM). **(E)** Cytokines and chemokine secretion in the medium of A375 cells was measured by multiplexed ELISA 24 hr after treatment with CDDP and MTX in the presence or absence of KIRA6 (1 μM). Data is expressed as log2 (fold change) of MFI values. In Western Blots β-actin was used as loading control. In all graphs values are presented as mean ± SD. Data is analyzed by one-way ANOVA except for one-sample t-test in (E),*p<0.05,**p<0.01.

**Figure 5 – figure supplement 1.**
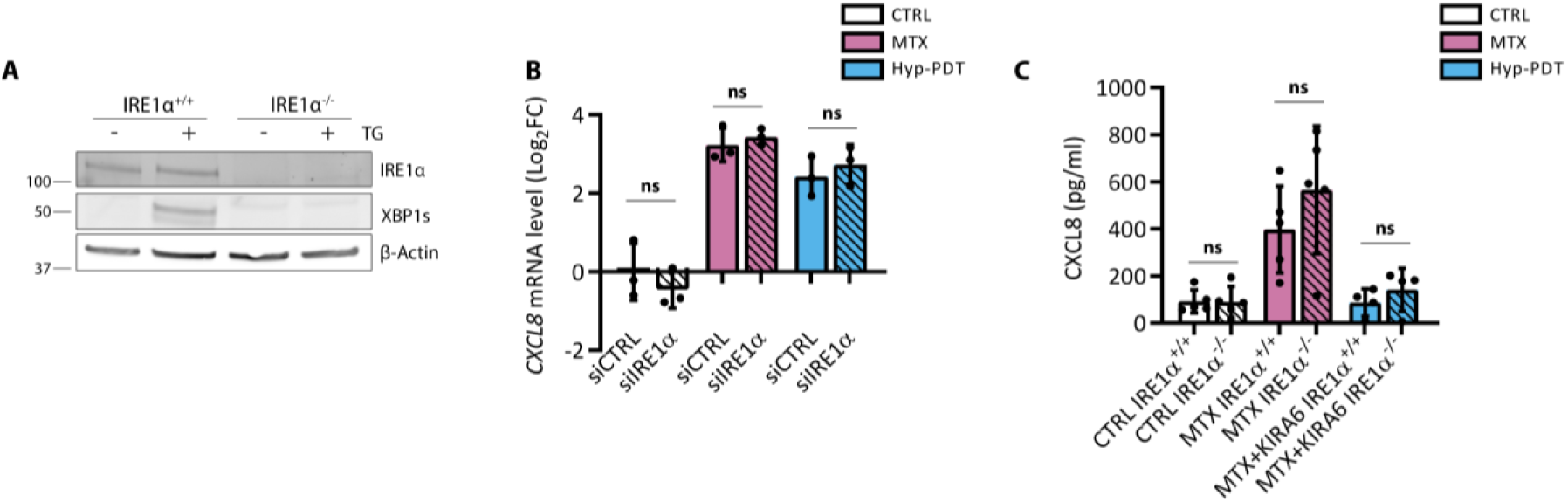
**(A)** Representative Western Blot showing the ability of thapsigargin (TG, 2 μM) to promote the splicing of XBP1 (XBP1s) 8 hr after treatment in IRE1α proficient and deficient A375 cells. β-actin was used as loading control. **(B)** The levels of CXCL8 mRNA induction were assessed by qPCR in A375 transfected with scrambled siRNA (siCTRL) or with IRE1α targeting siRNA (siIRE1α) in basal condition or 4 hr after treatment with MTX and Hyp-PDT. **(C)** CXCL1 secretion was measured by ELISA 24 hr after treatment with MTX in conditioned medium of murine embryonic fibroblasts (MEFs) proficient or deficient for IRE1α in the presence or absence of IRE1α inhibitor KIRA6 (1 μM). In all graphs values are presented as mean ± SD and analyzed by Two-way ANOVA, ns=not significant.

**Figure 6 – figure supplement 1.**
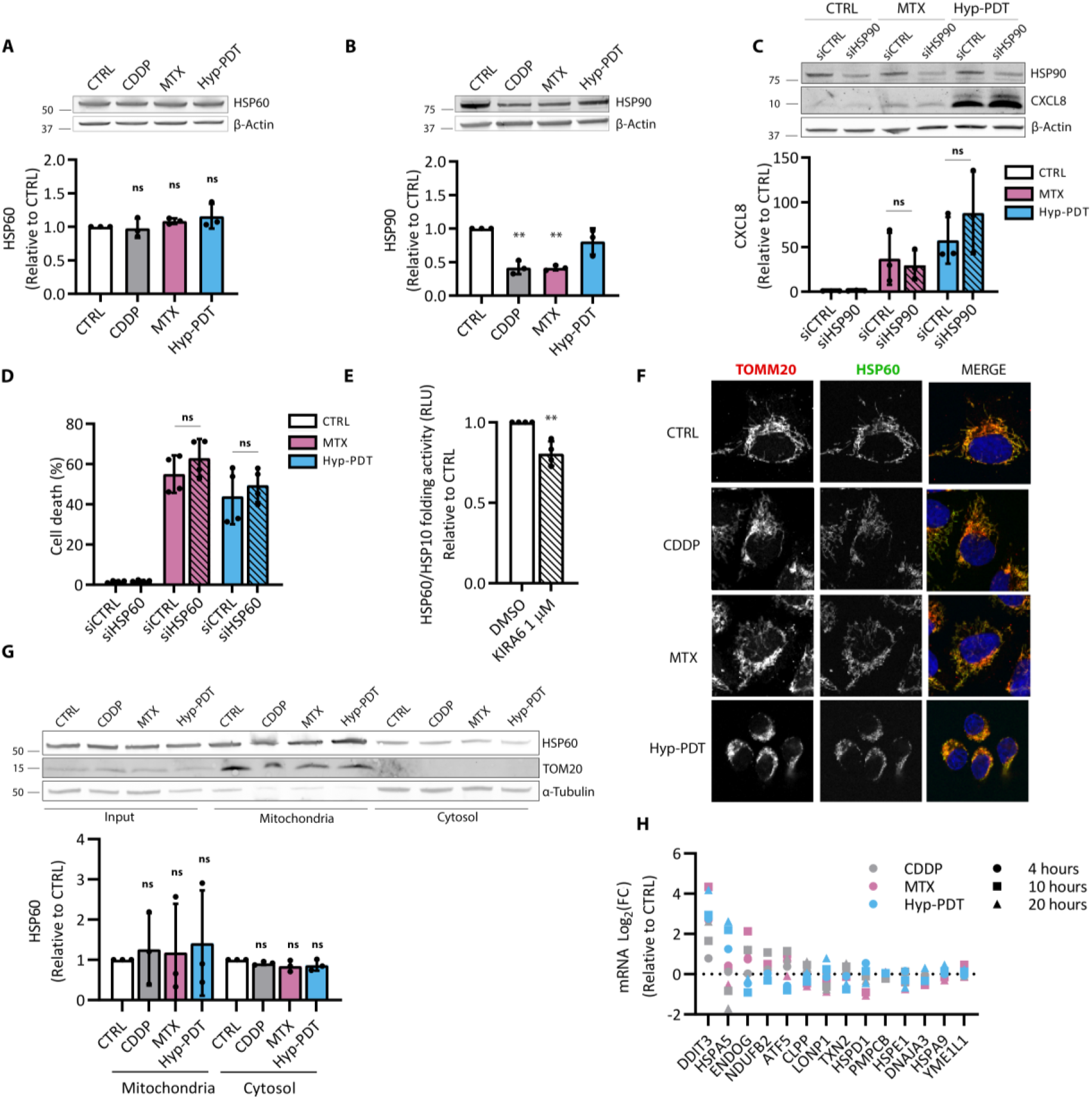
**(A-B)** Representative Western Blot and quantification of intracellular HSP60 and HSP90 content in A375 cells in basal conditions and 24 hr after treatment with CDDP, MTX and Hyp-PDT. **(C)** Representative Western Blot and quantification of the impact of HSP90 targeted siRNA (siHSP90) with respect to scramble siRNA (siCTRL) on intracellular CXCL8 accumulation 4 hr after treatment with MTX and Hyp-PDT. Data is expressed as fold change over control incubated with siCTRL. **(D)** Cell death was assessed in A375 cells 24 hr after treatment with CDDP, MTX and Hyp-PDT upon transfection with scramble siCTRL or siHSP60. **(E)** In vitro refolding activity of the HSP60/HSP10 chaperone complex after 1h of incubation with heat-mediated unfolded substrate proteins in control condition or in the presence of KIRA6 (10 μM). Data is expressed as fold change compared to control. **(F)** Representative confocal microscopy images for the intracellular localization of HSP60 (Green) and TOMM20 (red) in A375 cells 4 hr after treatment with CDDP, MTX and Hyp-PDT. Nuclei were counterstained with DAPI (blue). Scale bar: 10 μM. **(G)** Levels of HSP60 in the cytosol and mitochondria in basal condition or 4 hr after treatment with CDDP, MTX and Hyp-PDT were assessed by Western Blot after subcellular fractionation. TOMM20 was used as mitochondrial marker and α-tubulin was used as cytosolic marker. **(H)** Gene expression obtained from RNAseq data representing the log2(fold changes) of gens relative to the induction of mitochondrial UPR upon treatment with CDDP, MTX and Hyp-PDT compared to time-matched untreated control In all graphs values are presented as mean ± SD. Data is analyzed by one-sample t-test in (A,B,E,G) and Two-way ANOVA in (C,D), **p<0.01, ns= not significant.s

